# Detecting frequency-dependent selection through the effects of genotype similarity on fitness components

**DOI:** 10.1101/2022.08.10.502782

**Authors:** Yasuhiro Sato, Yuma Takahashi, Chongmeng Xu, Kentaro K. Shimizu

**Affiliations:** Department of Evolutionary Biology and Environmental Studies, University of Zurich, Winterthurerstrasse 190, 8057 Zurich, Switzerland; Research Institute for Food and Agriculture, Ryukoku University, Yokotani 1-5, Seta Oe-cho, Otsu, Shiga 520-2194, Japan; Graduate School of Science, Chiba University, Yayoi-cho 1-33, Inage-ku, Chiba 263-8522, Japan; Kihara Institute for Biological Research, Yokohama City University, Maioka 641-12, Totsuka-ward, Yokohama 244-0813, Japan

**Author notes:** Co-correspondence (email).

**Keywords:** Frequency-dependent selection, Genome-wide association study, Pairwise interaction model, Selection gradient analysis

## Abstract

Frequency-dependent selection (FDS) drives an evolutionary regime that maintains or disrupts polymorphisms. Despite the increasing feasibility of genetic association studies on fitness components, there are a few methods to uncover the loci underlying FDS. Based on a simplified model of pairwise genotype–genotype interactions, we propose a linear regression that can infer FDS from observed fitness. The key idea behind our method is the inclusion of genotype similarity as a pseudo-trait in selection gradient analysis. Single-locus analysis of *Arabidopsis* and damselfly data could detect known negative FDS on visible polymorphism that followed Mendelian inheritance with complete dominance. By extending the single-locus analysis to genome-wide association study (GWAS), our simulations showed that the regression coefficient of genotype similarity can distinguish negative or positive FDS without confounding other forms of balancing selection. Field GWAS of the branch number further revealed that negative FDS, rather than positive FDS, was enriched among the top-scoring single nucleotide polymorphisms (SNPs) in *Arabidopsis thaliana*. These results showed the wide applicability of our method for FDS on both visible polymorphism and genome-wide SNPs. Our study provides an effective method for selection gradient analysis to understand the maintenance or loss of polymorphism.

## 1 Introduction

Widespread polymorphism is a remarkable hallmark of the genomes of wild organisms. Balancing selection occurs when multiple alleles are maintained at a single locus through negative frequency-dependent selection (FDS), overdominance, and spatiotemporal variation in selection pressure (Hedrick, 2007). Among these mechanisms of balancing selection, negative FDS favors rare alleles over common alleles and consequently maintains multiple alleles at a locus. To date, negative FDS has been reported to act on various polymorphic traits such as coloration (Gigord *et al*., 2001; Takahashi *et al*., 2010; Le Rouzic *et al*., 2015; Nosil *et al*., 2018), self-incompatibility (Llaurens *et al*., 2008; Joly and Schoen, 2011; Shimizu and Tsuchimatsu, 2015), and resistance to natural enemies (Antonovics and Ellstrand, 1984; Brunet and Mundt, 2000; Sato and Kudoh, 2017). Conversely, positive FDS favors common alleles over rare alleles, thus unbalancing polymorphisms (Borer *et al*., 2010; Garrido *et al*., 2016).

High-throughput genotyping technology has now enabled us to evaluate the genomic basis of ecologically important traits and/or fitness in wild organisms (Durham *et al*., 2014; Fisher *et al*., 2016; Nosil *et al*., 2018; Exposito-Alonso *et al*., 2019; Tsuchimatsu *et al*., 2020). Specifically, genome-wide single nucleotide polymorphism (SNP) data help depict fitness-genotype association across genomes. In genome-wide association studies (GWASs) and quantitative trait locus (QTL) mapping, many statistical genetic analyses are based on linear regressions of a trait on genotype values (Broman and Sen, 2009; Gondro *et al*., 2013). If the target trait is a direct component of fitness, the regression coefficient of each locus describes the relationship between individual’s fitness and character — that is, selection gradient (Lande and Arnold, 1983) — at a target locus. By repeating this selection gradient analysis for all SNPs, for instance, Exposito-Alonso *et al*. (2019) quantified directional selection on a genome of *Arabidopsis thaliana*.

Despite its increasing appreciation in directional selection, little is known about the applicability of genetic association studies for FDS. To develop a general model of FDS, population genetics theory has long modelled genotype’s fitness as a product of the frequency of a focal genotype and its encounter frequency with the other genotypes, called pairwise interaction model (Schutz and Usanis, 1969; Cockerham *et al*., 1972; Asmussen and Basnayake, 1990; Trotter and Spencer, 2007; Schneider, 2008). Originally developed for direct competition among neighboring plants (Schutz and Usanis, 1969), the pairwise interaction model has thus far been introduced for direct competition and mating in insect populations (Álvarez-Castro and Alvarez, 2005). However, growing body of evidence suggests that FDS occurs not only through direct interactions but also through the indirect ones between genotypes (Antonovics and Ellstrand, 1984; Gigord *et al*., 2001; Takahashi *et al*., 2010; Sato and Kudoh, 2017). For instance, natural enemies mediate negative FDS on plant resistance when they attack undefended plants surrounding defended plants (Antonovics and Ellstrand, 1984; Brunet and Mundt, 2000; Sato and Kudoh, 2017). Mate choice in male insects also mediates negative FDS on female color polymorphism when male insects learn about different colors of female individuals (Van Gossum *et al*., 2001; Takahashi *et al*., 2010). Furthermore, while the pairwise interaction model assumes random interactions among individuals within subpopulations, e.g., mobile insects within split cages (Cosmidis *et al*., 1999; Fitzpatrick *et al*., 2007; Takahashi *et al*., 2014) or within split plant patches (Sato and Kudoh, 2017), the local action of FDS among neighboring plants is also plausible in a continuous vegetation (Janzen, 1970; CONNELL, 1971; Browne and Karubian, 2016) (also known as Janzen–Connell effects). If the pairwise interaction model can be exchanged with a general regression model, selection gradient analysis of FDS would become feasible in diverse organisms under various spatial structures (Fig. 1a).

**Figure 1:**
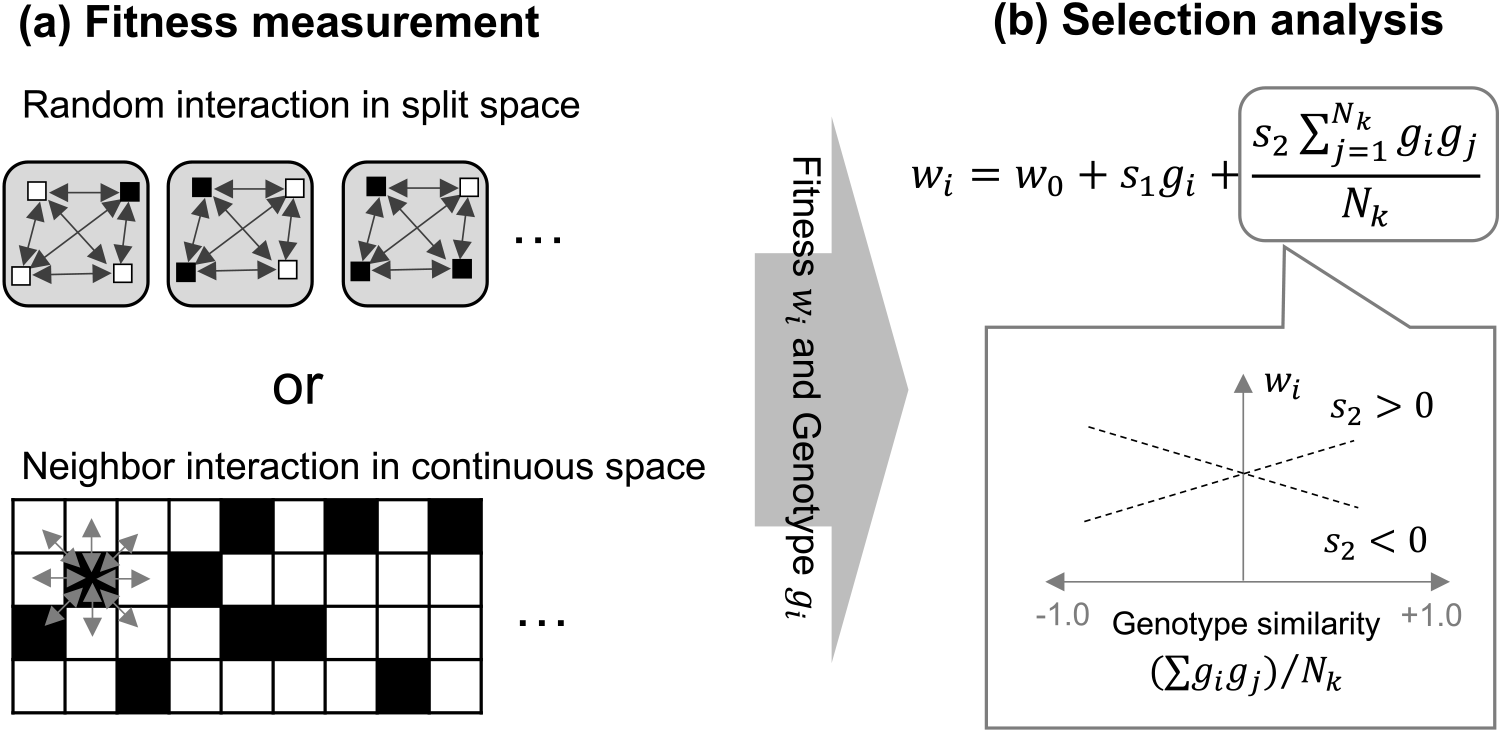
Presumable scheme from the fitness measurement (a) to selection gradient analysis (b). (a) Individual fitness is observed as a consequence of pairwise interactions (two-way arrows) among individuals carrying different genotypes (black and white squares) within split subpopulations (top gray squares: cf. the case of *A. halleri* and *I. elegans* in this study) or a continuous space (bottom: cf. the case of *A. thaliana*). (b) The selection gradient is then analyzed based on regression of the observed fitness *w*_*i*_ on genotypes *g*_*i*_. The second term in the equation is the same as that in Equation (1) and indicates how the second selection coefficient *s*_2_ represents the effects of genotype similarity between the genotypes *g*_*i*_ and *g*_*j*_.

Previously, we proposed regression models that incorporated neighbor genotype similarity into genetic association studies (Sato *et al*., 2021b,a). This idea was inspired by a model of ferromagnetism, which is widely known as the Ising model (Cipra, 1987). The pairwise physical interaction between two magnets that attract or repel each other may provide evolutionary insights into the effects of genetic interactions on spatiotemporal patterns and fitness consequences (Sato *et al*., 2021b). If two genotypes benefit from their similarity, these positive interactions force similar genotypes to be clustered within the local space. If two genotypes benefit from their dissimilarity, these negative interactions force the mixing of dissimilar genotypes across a space. Such a forward problem of the Ising model has been subject to genetic algorithms that mimic biological evolution (Anderson *et al*., 1991; PrÜgel-Bennett and Shapiro, 1997) because population optima are achieved through a series of updates on individual fitness. In contrast, an inverse problem of the Ising model deals with how to estimate the coefficient of physical interactions from individual energy that is analogous to individual fitness. These analogies between the Ising model and biological evolution led us to hypothesize that the direction and strength of pairwise genetic interactions could be estimated by incorporating genotype similarity as a pseudo-trait into selection gradient analyses (Fig. 1b).

In this study, we aimed to develop an effective selection analysis that infers FDS based on observed fitness. Specifically, we investigated whether the Ising-based model of pairwise genetic interaction (i) can accurately detect known FDS via single-locus analysis of visible polymorphic traits and (ii) screen FDS-associated polymorphisms from genome-wide SNPs. To accomplish these objectives, we first developed and applied the single-locus analysis to two examples under the split setting (Fig. 1a upper), including herbivore-mediated negative FDS on the trichome dimorphism of a wild herb *Arabidopsis halleri* (Sato and Kudoh, 2017) and male-mediated negative FDS on the female color polymorphism of a damselfly *Ischnura elegans* (Takahashi *et al*., 2014). The fact that trait expression of both trichome dimorphism and female color polymorphism followed Mendelian inheritance with complete dominance (Shimizu, 2002; SÁnchez-GuillÉn *et al*., 2005; Kawagoe *et al*., 2011) allowed us to substitute phenotype frequencies for genotype frequencies of the heterozygote and dominant homozygote. Extending the single-locus analysis to GWAS simulation, we then examined whether our method can aid in detecting simulated FDS from genome-wide SNPs. Distinct population structures were assumed in this GWAS simulation: The split setting exemplified mobile animals in split cages with variable morph frequencies (Takahashi *et al*., 2014) or plants in split plots (Sato and Kudoh, 2017), where individuals are expected to interact uniformly within the cage or plot (Fig. 1a upper). Contrastingly, the continuous setting exemplified the Janzen–Connell effects (Janzen, 1970; CONNELL, 1971) in a forest or grassland, in which sessile organisms interact only with their neighbors and FDS is restricted to local areas (Fig. 1a lower). The extended GWAS method was finally applied to the branch number data on *A. thaliana* accessions to screen polymorphisms associated with FDS under the continuous setting (Fig. 1a lower).

## 2 Methods

### 2.1 Model development

We developed a statistical method linking FDS, the Ising model, and a pairwise interaction model. First, we modeled pairwise interactions based on the forward problem of the Ising model. Then, we proposed a linear regression as an inverse problem of the Ising model. Finally, we examined fitness functions in a panmictic population to show how regression coefficients infer negative or positive FDS.

#### 2.1.1 Pairwise interactions based on the Ising model

To model pairwise interactions between genotypes, we focused on the Ising model of ferromagnetics. Let us assume that diploid organisms interact within panmictic subpopulations and produce their offspring based on the realized fitness (Fig. 1a top). We assume that a subpopulation *k* belongs to the meta-population *K* as *k* ∈ *K*, where two individuals *i* and *j* belong to subpopulation *k* such that *i, j* ∈ *k*. We further assumed that individuals *i* and *j* had two alleles at a locus, with the ancestral allele A being dominant over the derived allele a, where the genotype values were encoded as *g*_*i*(*j*)_ ∈ {AA, Aa, aa} = {+1, +1, -1}. This dominant encoding represented well-reported cases in which FDS often acts on a trait exhibiting dimorphism with complete dominance (e.g., Takahashi *et al*., 2010; Sato and Kudoh, 2017). By incorporating the genotype similarity between *i* and *j*, we designated fitness *w* for individual *i* as follows:

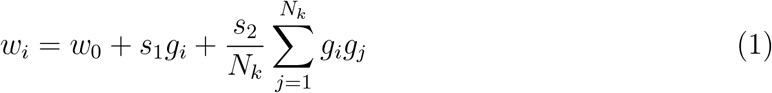

where *w*_0_ indicates the base fitness; *s*_1_ and *s*_2_ indicate the selection coefficients for self-genotype effects and genotype similarity, respectively; and *N*_*k*_ indicates the total number of individuals within subpopulation *k*. The genotype similarity 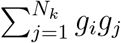 represents the similarity (or difference) of the genotype composition of the subpopulation from the individual *i*. If two individuals share the *same* genotype values, *g*_*i*_*g*_*j*_ = (+1) × (+1) = (−1) × (−1) = +1. If two individuals have *different* genotype values, *g*_*i*_*g*_*j*_ = (−1) × (+1) = (+1) × (−1) = −1. Thus, the within-population genotype similarity 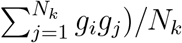 ranges from -1.0 (perfect dissimilarity) to +1.0 (perfect similarity; Fig. 1b) as scaled by the total number of interacting individuals within the subpopulation *N*_*k*_. Additionally, we could also assume an additive expression for a trait responsible for FDS as *g*_*i*(*j*)_ ∈ {AA, Aa, aa} = {+1, 0, -1}, where the products involving heterozygotes were considered 0 × 0 = 0 (neither similar nor dissimilar). This additive encoding enabled us to assume the intermediate strength of the FDS for an intermediate morph. However, most empirical studies have reported FDS between two out of multiple morphs (e.g., Gigord *et al*., 2001; Takahashi *et al*., 2010; Le Rouzic *et al*., 2015; Sato and Kudoh, 2017; Nosil *et al*., 2018). As intermediate FDS on an additive trait is still uncommon, in this study, we focussed on dominant encoding.

Analogous to the Ising model, the forward problem of Equation (1) is to optimize *w*_*i*_ with given *s*_1_ and *s*_2_ by modulating *g*_*i*(*j*)_. This was analogous to biological evolution, where *s*_1_ and *s*_2_ selected genotypes *g*_*i*(*j*)_ based on the fitness *w*_*i*_. The point of this forward problem of the Ising model is that negative or positive *s*_2_ favored mixed (i.e., locally negative FDS) or clustered (locally positive FDS) genotype distributions in a lattice space, respectively (Figure S1a,b). The stochastic simulation based on Equation (1) is given in Appendix S1, Table S1, and Figure S1.

#### 2.1.2 Regression model as an inverse problem of the Ising model

To infer FDS from the inverse problem of the Ising model, we modified Equation (1) as a regression model. We redefine the individual fitness *w*_*i*_ as the response variable *y*_*i*_; the genotype *g*_*i*(*j*)_ as the explanatory variables *x*_*i*(*j*)_; the base fitness *w*_0_ as the intercept *β*_0_; and the selection coefficients *s*_1_ and *s*_2_ as the regression coefficients *β*_1_ and *β*_2_, respectively. We also added a residual error *e*_*i*_ to Equation (1) to obtain a statistical model as follows:

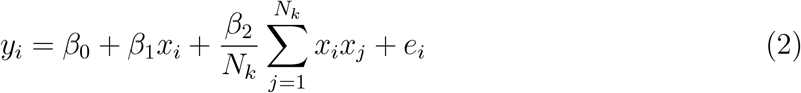

where Equation (2) poses a regression analysis to estimate 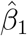 and 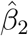 from the given *y*_*i*_ and *x*_*i*(*j*)_. According to the inference from *s*_2_ (Appendix S1), the negative or positive *β*_2_ represents a negative or positive FDS between two alleles, respectively.

When *y*_*i*_ is an absolute fitness, the FDS may act asymmetrically between the two alleles. For example, negative FDS on relative fitness is known to occur in the Hawk-Dove game (Takahashi *et al*., 2018), where hawks profit from competition with doves, whereas doves suffer from competition with hawks. Thus, for the correct inference of FDS, it was necessary to consider asymmetric and symmetric FDS. Asymmetric FDS can be described by a multiplicative interaction between the second and third terms of Equation (2) (Sato *et al*., 2021b) and is expressed as follows:

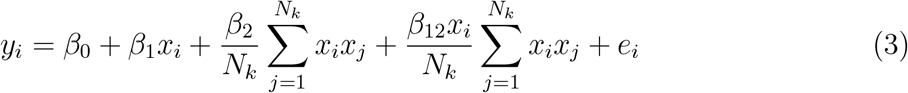

where *β*_12_ indicates the coefficient for the asymmetric effects of genotype similarity. If *β*_12_ was statistically significant, the slope coefficient of the genotype similarity differed among focal genotypes (Sato *et al*., 2021b), meaning that the strength or direction of FDS is asymmetric between genotypes. When *y*_*i*_ is a relative fitness, negative fitness effects on one allele coincide with positive effects on another allele, and asymmetric FDS would be unnecessary. By extending Equations (2) and (3) to mixed models, we could also apply these regression methods for GWAS (Appendix S2).

#### 2.1.3 Fitness functions with respect to regression coefficients

To clarify how the coefficients for genotype similarity effects *β*_2_ and asymmetric effects *β*_12_ corresponded to FDS, we finally analyzed Equation (3) as a function of allele frequencies. We suppose that all individuals uniformly interact in a sufficiently large population with random mating (i.e., *N*_*k*_ → ∞). The likelihood of one genotype interacting with the other genotypes depends on genotype frequencies derived from an allele frequency within a population (Appendix S3). Let *f* be the frequency of the A allele within the panmictic infinite population. The ratio of genotype frequency on panmixia was as follows: AA: Aa: aa = *f* ^2^ : 2*f* (1 − *f*) : (1 − *f*)^2^. Assuming the complete dominance of the A allele over the a allele (*x*_*i*(*j*)_ ∈ {AA, Aa, aa} = {+1, +1, -1}), we calculated all the combinations among the three genotypes (Table S2) and consequent fitness *y*_*i*_ for AA, Aa, and aa genotypes as

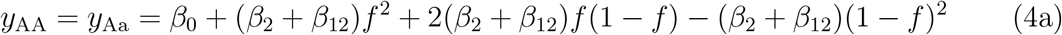

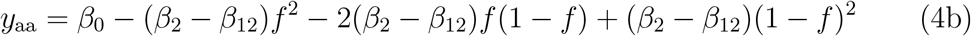

where *y*_AA_, *y*_Aa_, and *y*_aa_ denote the fitness values for the AA, Aa, and aa genotypes, respectively (Appendix S3). The allele-level marginal fitness was then defined by weighting the genotype fitness with the allele frequency as follows:

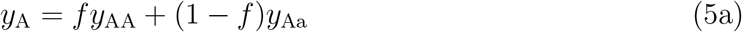

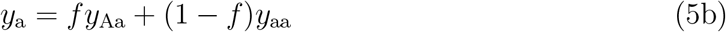

Figure 2 shows how the fitness value Equations (5a) and (5b) vary in response to the allele frequency *f*. Symmetric negative FDS was exemplified by the negative *β*_2_ without any asymmetric effects *β*_12_ (i.e., *β*_12_ = 0; Fig. 2a). Asymmetric negative FDS was described by the negative (Fig. 2c) or positive (Fig. 2e) asymmetric effect *β*_12_, where the negative *β*_2_ denoted negative FDS on the relative fitness between two alleles (Fig. 2c, e). In contrast, symmetric positive FDS was exemplified by the positive *β*_2_ with no (Fig. 2b), negative (Fig. 2d), or positive (Fig. 2f) values of the asymmetric effect *β*_12_. In summary, the sign of the symmetric effects *β*_2_ represents negative or positive FDS on relative fitness between two alleles even when the asymmetric effects *β*_12_ are not zero.

**Figure 2:**
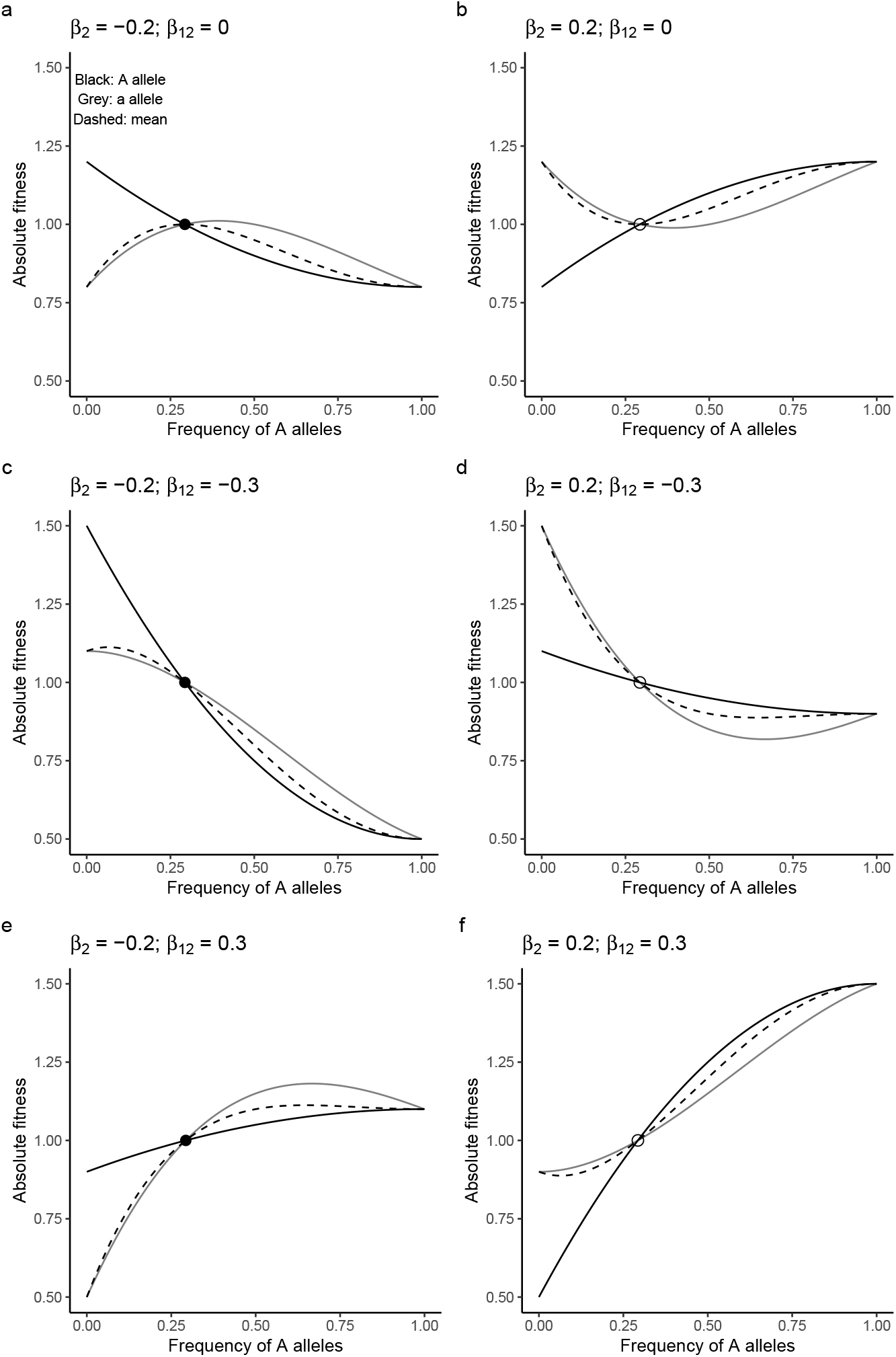
Numerical examples for the fitness values *y*_*i*_ in response to allele frequency when the A allele is completely dominant over the a allele. The black and gray lines indicate the marginal fitness of A and a alleles; that is, Equations (5a) and (5b), respectively. Dashed curves indicate the population-level mean fitness between the two alleles Equation (6). (a) Symmetric negative FDS, (b) symmetric positive FDS, (c and e) asymmetric negative FDS, and (d and f) asymmetric positive FDS. Closed and open circles indicate stable or unstable states, respectively. The base fitness and no directional selection were set at *β*_0_ = 1.0 and *β*_1_ = 0.0 for all panels.

While the Ising model poses an optimization problem of the total energy based on its interaction coefficient (Cipra, 1987; Anderson *et al*., 1991; PrÜgel-Bennett and Shapiro, 1997), evolutionary biologists have long analyzed optima of population-level mean fitness under FDS (Cockerham *et al*., 1972; Schneider, 2008; Takahashi *et al*., 2018). In a panmictic population, population-level mean fitness is given by the allele-level marginal fitness weighted by its allele frequency (Appendix S3); that is, the weighted mean is given as follows:

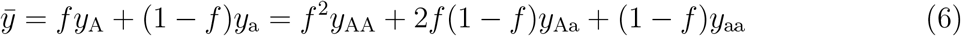

This population-level mean fitness is maximized at an intermediate frequency under symmetric negative FDS (Fig. 2a) (Schneider, 2008). This expectation from randomly interacting and mating populations is comparable to the forward problem of the Ising model when *s*_2_ *<* 0, where spatially mixed genotypes increase the population sum of *y*_*i*_ more than monomorphic populations (Sato *et al*., 2021b). Even under an asymmetric negative FDS (Fig. 2c and e), the population-level mean fitness became larger than expected by a weighted mean of monomorphic populations (= frequency-independent selection) (Takahashi *et al*., 2018) (Fig. 2c and e). In contrast, the population-level mean fitness was minimized at an intermediate frequency under a symmetric positive FDS (Fig. 2b). Under asymmetric positive FDS, the population-level mean fitness was neither maximized nor greater than expected by frequency-independent selection (Schneider, 2008; Takahashi *et al*., 2018) (Fig. 2d and f). When A and a alleles show additive expression as *x*_*i*(*j*)_ ∈ {AA, Aa, aa} = {+1, 0, -1}, fitness functions become so complicated that more than two equilibria might arise (Appendix S3; Figure S2) but have rarely been reported empirically.

### 2.2. Single-locus examples

To test whether the known FDS could be detected using our method, we applied the single-locus analysis equations (2) and (3) to *Arabidopsis* and *Ischnura* data (Sato and Kudoh, 2017; Takahashi *et al*., 2014) collected under a split setting (Fig. 1a, top). The fitness components in the real data were absolute values (e.g., number of eggs, flowers, and reproductive branches) where asymmetric FDS on the absolute fitness (Fig. 2c-d) was possible in addition to symmetric FDS on relative fitness (Fig. 2a and b). Therefore, symmetric and asymmetric FDS were tested using Equations (2) and (3), respectively. The lme4 package (Bates *et al*., 2015) in R was used for the single-locus analyses.

#### 2.2.1 Flower production of hairy and glabrous plants

We applied single-locus analysis for the flower production data of *Arabidopsis halleri* subsp. *gemmifera* to detect known negative FDS mediated by leaf beetle attacks on hairy and glabrous plants (Sato and Kudoh, 2017). The original data of Sato and Kudoh (2017) are downloaded from the Dryad repository (https://doi.org/10.5061/dryad.53k2d) and were re-analyzed using our proposed method. Sato and Kudoh (2017) set circular split plots (1 m in diameter) and recorded the trichome phenotype (hairy or glabrous), the number of flowers, leaf damage score, and the length of largest leaf for all individual plants within each plot. Field surveys were conducted along a 200-m line transect at the Omoide River, Hyogo, Japan (35°06′N, 134°56′E), from 2013 to 2016. The total sample size was 3,070 individuals among 324 plots. According to Sato and Kudoh (2017), we used a generalized linear mixed model (GLMM) with a Poisson error structure and a log-link function. The response variable was the number of flowers. The fixed effects were the self-phenotype (hairy or glabrous), similarity between the two morphs, and total number of plants within each field plot. The log-transformed length of the largest leaf (mm), which reflects the plant size, was included as an offset term. The random effects were the field plot IDs nested below the study years. In *A. halleri*, hairy alleles are known to be dominant over glabrous alleles at the *GLABRA1* locus (Shimizu, 2002; Kawagoe *et al*., 2011). Given the complete dominance of hairy alleles, we assumed complete dominance at the *GLABRA1* locus with *x*_*i*(*j*)_ ∈ {AA, Aa, aa} = {+1, +1, -1} on the basis of the trichome phenotype of an individual.

#### 2.2.2 Egg production of andromorph and gynomorph females in a damselfly

We also applied single-locus analysis for the egg production by the blue-tailed damselfly *Ischnura elegans* to detect known negative FDS and the consequent increase in population-level mean fitness between an andromorph and a gynomorph (Takahashi *et al*., 2014; Le Rouzic *et al*., 2015). The original data were derived from Takahashi *et al*. (2014) and consisted of 102 andromorphs and 79 *infuscans*-type gynomorphs. Takahashi *et al*. (2014) assigned adult *I. elegans* with an andromorph frequency of 0.2, 0.5, or 0.8 into split-field cages under low- or high-density conditions. The field experiment was conducted at the Stensoffa Field Station of Lund University. We used a GLMM with a Poisson error structure and a log-link function according to Takahashi *et al*. (2014). The response variable was the number of mature eggs. The fixed effects were morph type and morph similarity within a cage. The cage ID and experimental ID were considered random effects, where the cage ID was nested below the experimental ID. The andromorph allele is known to be dominant over the *infuscans*-type gynomorph allele on an autosomal locus (SÁnchez-GuillÉn *et al*., 2005). Therefore, on the basis of phenotype frequencies within the split cages, we assumed complete dominance with genotype values encoded as *x*_*i*(*j*)_ ∈ {AA, Aa, aa} = {+1, +1, - 1} for homozygous andromorphs, heterozygous andromorphs, and homozygous gynomorphs, respectively. The interaction term between the morph type and similarity was also considered in the line of GLMMs to test the significance of asymmetric FDS between the two morphs. If the interaction term was not significant, the coefficients of the main effect were estimated using GLMM without any interaction terms.

### 2.3 GWAS simulation

Simulations were used to evaluate the power of our method in detecting causal polymorphisms from genome-wide SNPs. The entire procedure consisted of three steps: We (i) simulated genomic structure under balancing selection, (ii) conducted virtual experiments to simulate fitness from the simulated genomes, and (iii) applied our method for GWAS of simulated fitness and genomes. We used Equation (2) to focus on symmetric FDS on relative fitness in this simulation because selection acted not on absolute but on relative fitness. To test the notion that linear mixed models (LMMs) usually outperform standard linear models (LMs) in GWAS (Kang *et al*., 2008), we compared the performance of both LMs [Equation (2)] and LMMs (see Appendix S2 for the mixed model extension). We used SLiM version 3 (Haller and Messer, 2019) for population genetic simulations; and the vcfR (Knaus and GrÜnwald, 2017), gaston (Perdry and Dandine-Roulland, 2020), rNeighborGWAS (Sato *et al*., 2021b), and pROC (Robin *et al*., 2011) packages in R version 4.0.3 (R Core Team, 2019) for GWAS mapping.

#### 2.3.1 Simulated genomes

Population genetic simulations were performed using SLiM version 3 to create a realistic genome structure. By running 30 independent iterations for 2000 generations, we simulated 50 kb nucleotide × three chromosomes × 10 subpopulations × 200 individuals. Base parameters were set as follows: mutation rate *μ* = 10^−6^, selection coefficient *s*_1_ = *s*_2_ = 0.1 for non-neutral mutations, and recombination rate *r* = 10^−5^. Ten subpopulations were distributed in a circle with a low migration rate *m* = 10^−4^ between neighboring populations. In this simulation, we decomposed the total fitness as *w*_*i*_ = *w*_*i*,1_ + *w*_*i*,2_. The first fitness component *w*_*i*,1_ involves self-genotype effects *β*_1_, and *w*_*i*,2_ is the second fitness component subject to the power analysis of genotype similarity effects *β*_2_. For self-genotype effects *β*_1_, we define stabilizing selection as 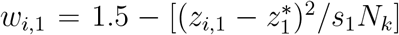, where 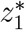 is the optimum number of QTLs responsible for self-genotype effects per genome per population and *z*_*i*_ is the number of QTLs for individual *i*. We set 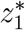 to 5 and assumed additive effects by the QTLs. Following the standard way to simulate polygenic selection (Haller and Messer, 2019), we did not substitute QTLs for *w*_1,*i*_ even after they were fixed. For the genotype similarity effects *β*_2_, we assumed four specific scenarios of selection on the second fitness component *w*_*i*,2_ as follows:

Scenario 1. Negative frequency-dependent selection: For the test of *β*_2_, we simulated negative FDS on the second fitness component *w*_*i*,2_. We simulated negative FDS for the second fitness component as *w*_*i*,2_ = 1.5 − *s*_2_*g*_*i*_*f*_*k*_, where *f*_*k*_ indicates the frequency of the mutation within a subpopulation *k*. Novel mutations were assumed to be recessive to ancestral alleles with genotype *g*_*i*_ redefined as *g*_*i*_ ∈ {AA, Aa, aa} = {1, 1, 0}. Polymorphisms were likely balanced under negative FDS; thus, the mutation rate was set at half of the base parameter to maintain the number of causal SNPs in the same order as that in the other scenario.

Scenario 2. Positive frequency-dependent selection: We also simulated the opposite regime, positive FDS, on the second fitness component *w*_*i*,2_. It is known that locally acting positive FDS within a subpopulation can lead to global coexistence of two alleles among subpopulations (Molofsky *et al*., 2001). Therefore, we separated the entire population into four panmictic subpopulations to simulate polymorphic loci. Similar to negative FDS, we simulated positive FDS as *w*_*i*,2_ = 1.5 + *s*_2_*g*_*i*_*f*_*k*_. Novel mutations were assumed to be recessive to ancestral alleles with genotype *g*_*i*_ redefined as *g*_*i*_ ∈ {AA, Aa, aa} = {1, 1, 0}.

Scenario 3. Overdominance: To test whether *β*_2_ confounded frequency-independent types of balancing selection, we simulated genomes under overdominance selection. The second fitness component is defined as *w*_*i*,2_ = 1.5 + *s*_2_*hg*_*i*_, where *h* is the dominance coefficient expressed on the basis of genotypes as {*h*_AA_, *h*_Aa_, *h*_aa_} = {1.0, 2.0, 1.0}.

Scenario 4. Spatiotemporally varying selection: To test another type of frequency-independent balancing selection, we simulated genomes under spatiotemporally varying selection. The second fitness component is defined as *w*_*i*,2_ = 1.5 + *sg*_*i*_, where *s* varies in space and time. We assumed *s*_2_ = 0.1 for two subpopulations, and *s*_2_ = −0.1 for the other two subpopulations. For six of the four subpopulations, we changed the selection coefficient in time to *s*_2_ = 0.1 for odd generations and *s*_2_ = −0.1 for even generations. We reset *m* = 0.001 to allow a higher migration rate. In this scenario, novel mutations were assumed to be recessive to ancestral alleles.

#### 2.3.2 Virtual experiments

Then, we sampled the simulated genomes and generated fitness values from the genotype data. The simulated genomes were exported in variant call format (.vcf) and loaded into R using the vcfR package (Knaus and GrÜnwald, 2017). SNPs were filtered with a cutoff threshold of a minor allele frequency (MAF) of 0.01. We assumed two experimental settings: split and continuous populations (Fig. 1a). To describe the two distinct cases, 900 individuals were randomly sampled from each simulation without replacement and assigned to a 30 × 30 lattice space for the continuous setting, or 10 individuals each to 90 split cages for the split setting.

Fitness values were then simulated from the simulated genomes and their spatial arrangement. To calculate the self-genotype component *β*_1_*x*_*i*_ in Equation (2), we assigned 0.1 (which corresponded to the selection coefficient *s*_1_ in the population genetic simulation) to *β*_1_ of causal SNPs or zero to those of the other SNPs. The second fitness component at causal SNPs was generated with *β*_2_ = 0.1 (corresponding to the strength of balancing selection *s*_2_ in the population genetic simulation) for negative or positive FDS as 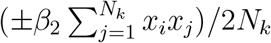, for overdominance as *x*_*i*_ ∈ {AA, Aa, aa} = {1 + *β*_2_, 1 + *β*_2_*h*, 1}, and for spatiotemporally varying selection as a random assignment of ±*β*_2_ to *β*_2_*x*_*i*_. Fitness variance was not adjusted for the first and second fitness components since the number of causal SNPs and their effect sizes were controlled during the population genetic simulation above. Gaussian residual errors were finally added to the simulated fitness such that approximately one-third of the total phenotypic variation was attributed to the environmental variance as Var(**e**) = (0.75)^2^ × Var(**w**). This range of the number of causal SNPs and the proportion of phenotypic variation explained by the model were based on parameter settings where the model performance was well differentiated (Sato *et al*., 2021b).

#### 2.3.3 GWAS using simulated genomes and fitness

Finally, we performed association mapping of the simulated fitness with respect to *β*_1_ and *β*_2_. The rNeighborGWAS package (Sato *et al*., 2021b) was used to implement the regression model in Equation (2) as an LMM (Appendix S2). The false vs. true positive rate was analyzed using the receiver operating characteristic (ROC) curve. Similar to the generative model, we assumed the dominant encoding for the three genotypes, *x*_*i*(*j*)_ ∈ {AA, Aa, aa} = {+1, +1, -1}. To measure the efficiency of causal polymorphism detection, we calculated the area under the ROC curve (AUC) for the -log_10_(*p*-values) of *β*_1_ or *β*_2_. The AUC ranged from 0.5 (no power to detect causal SNPs) to 1.0 (perfect matching between the top *p*-value score and causal SNPs). To measure the accuracy of the effect size estimates, we compared the true and estimated values of *β*_2_. To test whether LMMs could outperform standard LMs, we also compared AUCs and 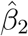 between LMM and LM.

### 2.4 Pilot GWAS

To examine whether our method is applicable to the real GWAS dataset, we conducted a pilot GWAS of the reproductive branch number in field-grown *A. thaliana* under a continuous setting (Fig. 1a lower). According to Sato *et al*. (2019), a summer cohort composed of natural accessions with various life cycles was established to investigate the survival and reproduction under stressful environments. We selected 199 worldwide accessions from 2029 inbred lines sequenced by the RegMap (Horton *et al*., 2012) and 1001 Genomes project (Alonso-Blanco *et al*., 2016). Full-imputed genotypes were downloaded from the AraG-WAS catalog (Togninalli *et al*., 2018). For the 199 accessions, 1,819,577 SNPs were selected at a cut-off threshold of MAF *>* 0.05. Three replicates of the 199 accessions were sown on Jiffy-seven (33 mm in diameter) and stratified under constant dark conditions at a temperature of 4C° for 1 week. Seedlings were first grown under short-day conditions (8 L: 16D, 20C°) for 6 weeks. Individual plants were then potted into a plastic pot (6 cm in diameter) filled with mixed soils of agricultural compost (Profi Substrat Classic CL ED73, Einheitserde Co.) and perlite with a 3:1 L ratio of perlite. Potted plants were transferred to the common garden at the Irchel Campus of the University of Zurich (Zurich, Switzerland: 47°23′N, 08°33′E) on 8 July 2019. In the field setting, a set of 199 accessions and an additional Col-0 accession were randomly assigned to each block without replacement. The 200 plants were set in plastic trays (10 × 40 cells in a continuous space) in a checkered pattern. Three replicates of each block were set *>* 1.5 m apart from each other. We recorded the length of the largest leaf (mm) at the beginning of the experiment, the presence of bolting after 2 weeks, and the branch number at the end of the experiment (27 August 2019). We considered the branch number as a proxy for fitness because it is known as a major fitness component of *A. thaliana* (Chong *et al*., 2018) and other reproductive phenotypes were difficult to observe owing to the stressful summer environment. Dead plants were recorded as a branch number of zero, *i*.*e*., with no fitness. Accession names and phenotype data are presented in Table S3.

The branch number was analyzed as a target trait of GWAS. The rNeighborGWAS package version 1.2.3 (Sato *et al*., 2021b) was used to implement Equations (2) and (3) as GWAS (Appendix S2). The inbred lines of *A. thaliana* have either AA or aa genotype, in which the qualitative interpretation of *β*_2_ and *β*_12_ in this inbred case remains the same as in the case of complete dominance (Appendix S3; Figure S3). The response was log(x+1)-transformed number of branches. We assumed that FDS arose from genetic interactions among neighboring plants in small *Arabidopsis*, and thus, the genotype similarity was considered up to the nearest neighbors; that is, *N*_*k*_ = 4 from a focal individual. The initial plant size, presence of bolting, experimental block ID, and edge of each plot (or not) were considered as non-genetic covariates. The marker kinship and genome-wide structure of neighbor genotype similarity were considered random effects. After association mapping, we focused on SNPs with a -log_10_(*p*-value) score *>* 4.0. We searched candidate genes within ∼10 kb around the target SNPs based on the Araport11 gene model with the annotation of The Arabidopsis Information Resource (TAIR; accessed on 31 December 2021). Gene ontology (GO) enrichment analysis was conducted using the Gowinda algorithm (Kofler and SchlÖtterer, 2012) with the options “–gene-definition undownstream1000,” “–min-genes 2,” and “–mode gene.” The GO.db package (Carlson, 2020) and the latest TAIR AGI code annotations were used to build the input files.

## 3 Results

### 3.1 Negative FDS on trichome dimorphism in *A. halleri*

To test whether the single-locus analysis could detect the known negative FDS, we applied Poisson GLMM for the flower production data on hairy and glabrous plants of *A. halleri* under the split setting of field plots (Sato and Kudoh, 2017). The Poisson GLMM detected a negative and significant coefficient of the morph similarity 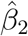 (Table 1a), indicating a negative FDS where a focal plant produced more flowers as dissimilar morphs were grown within the same plot. The lack of significance of the interaction term between the trichomes and morph similarity suggest that negative FDS is symmetric between hairy and glabrous plants (Table 1a). We also found the same level of the self-morph coefficient 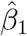 as the negative coefficient of morph similarity 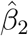 (Table 1a), which indicates the simultaneous action of directional selection and negative FDS between the two morphs. However, the total number of plants, namely, the density, had no significant effect on flower production (Table 1a). The fitness function estimated from 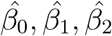, and 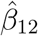 showed that a stable equilibrium under negative FDS remained at a biased but not monomorphic frequency (Fig. 3a). This discrepancy between the observed frequency and expected equilibrium was likely because the present selection analysis could not incorporate another major component of fitness in *A. halleri that is* clonal reproduction (Sato and Kudoh, 2017). These results provide qualitative evidence for negative FDS on trichome dimorphism through the fitness component of flower production.

**Table 1:** Poisson generalized linear mixed model (GLMM) applied to the number of flowers between hairy and glabrous *A. halleri* (a) or the number of mature eggs between the andromorph and gynomorph of *I. elegans* (b). Morph similarity was calculated from the genotype similarity defined by Equation (2). Estimated coefficients, their standard errors (SE), *Z*-values, and *p*-values are shown for multiple regressions. Bold letters indicate significance at *p <* 0.05 by Wald tests.

| Fixed effects | Coefficient | SE | $Z$ | $p$ |
| --- | --- | --- | --- | --- |
| **Intercept** $\hat{\beta}_0$ | **-2.076** | **0.17** | **-11.97** | **<2e-16** |
| **Self-morph** $\hat{\beta}_1$ | **0.146** | **0.008** | **17.32** | **<2e-16** |
| **Morph similarity** $\hat{\beta}_2$ | **-0.163** | **0.018** | **-9.22** | **<2e-16** |
| Total no. of plants | -0.006 | 0.010 | -0.63 | 0.53 |
| Self × Similarity $\hat{\beta}_{12}$ | -0.088 | 0.048 | -1.81 | 0.07 |

(b) *Ischnura elegans*
| Fixed effects | Coefficient | SE | $Z$ | $p$ |
| --- | --- | --- | --- | --- |
| **Intercept** $\hat{\beta}_0$ | **4.55** | **0.191** | **23.8** | **<2e-16** |
| **Self-morph** $\hat{\beta}_1$ | **-0.021** | **0.008** | **-2.56** | **0.01** |
| **Morph similarity** $\hat{\beta}_2$ | **-0.878** | **0.022** | **-39.30** | **<2e-16** |
| Density | 0.097 | 0.237 | 0.41 | 0.68 |
| Self × Similarity $\hat{\beta}_{12}$ | -0.042 | 0.25 | -0.17 | 0.87 |

**Figure 3:**
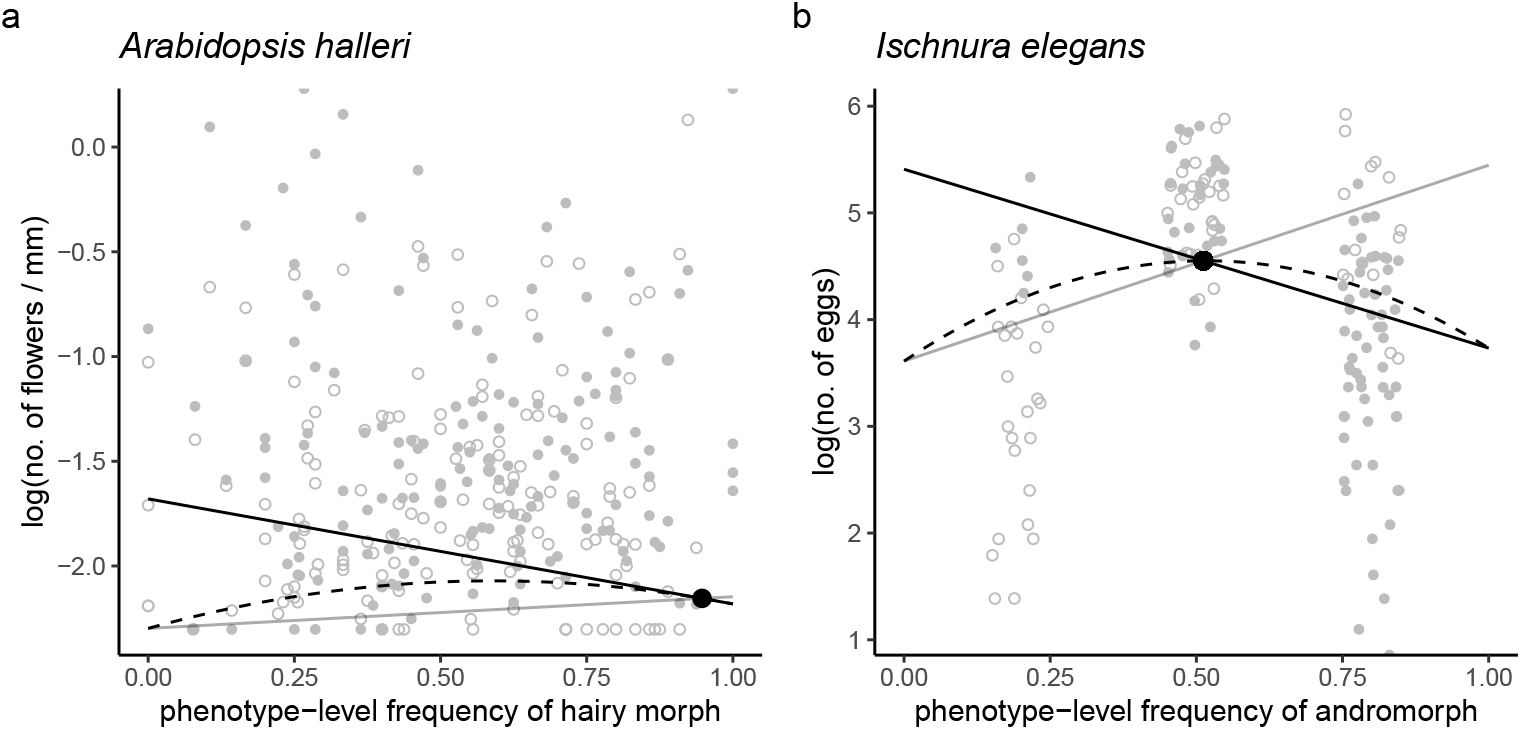
Negative frequency dependence in (a) the size-adjusted number of flowers in *A. halleri* and (b) the number of mature eggs in *I. elegans*. Fitness functions are based on the estimates from Table 1 and are shown at a phenotype level (see case 3 in Appendix S3) to project model trends on the observed fitness. The black and grey lines indicate the fitness function of dominant (i.e., hairy and andromorph) or recessive (glabrous and gynomorph) morphs, respectively. Filled and open circles indicate the observed fitness for the dominant or recessive morph within a plot, respectively. The dashed curve shows the population-level mean fitness. The black dot represents a stable equilibrium under negative FDS. In panel (a), a single circle corresponds to a field plot where the average number of flowers adjusted by plant size (mm) among hairy or glabrous plants is given after log(*x*+0.1)-transformed.

### 3.2 Negative FDS on female color polymorphisms in *I. elegans*

To further test whether the single-locus analysis could represent a known relationship between negative FDS and population-level mean fitness, we applied Poisson GLMM for the data on the number of mature eggs between the andromorph and gynomorph of *I. elegans* under the split setting of field cages (Takahashi *et al*., 2014). Consistent with the previous evidence for negative FDS (Van Gossum *et al*., 2001; Le Rouzic *et al*., 2015), we found a significantly negative coefficient of morph similarity 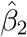 (Table 1b). The interaction term between the morph type and similarity was not significant (Table 1b), indicating no significant asymmetry in the negative FDS between the two morphs. We also found a significant effect of the morph type (andromorph or gynomorph) on the number of eggs, but its effect was much less significant than that of morph similarity (Table 1b). As reported in a previous study (Takahashi *et al*., 2014), the density did not significantly affect the egg number (Table 1b). The fitness function estimated from 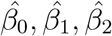, and 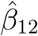 shows that negative FDS allows for the coexistence of the two morphs at an intermediate frequency (Fig. 3b). Consistent with the results of Takahashi *et al*. (2014), the population-level mean fitness increased at a stable equilibrium at the intermediate frequency (Fig. 3b). These results support the action of symmetric negative FDS and the consequent increase in the population-level mean fitness. Taken together, *A. halleri* and *I. elegans* data suggest that our single-locus model can be applied to the selection gradient analyses of FDS on visible polymorphic traits that follow typical Mendelian inheritance with complete dominance (see also the discussion “Selection gradient along genotype similarity”).

### 3.3 Detection of simulated FDS across a genome

We simulated genotypes and fitness to test whether our method could distinguish negative FDS, positive FDS, overdominance, and spatiotemporally varying selection among genomewide SNPs. The simulated genomes had 2,000 to 4,500 SNPs with MAFs *>* 0.01 across 50 kbp nucleotide sequences (Fig. S4). They exhibited low heterozygosity (*H*_t_ *<* 0.1) and moderate to strong differentiation among 10 populations (*G*_st_ *<* 0.5; Fig. S4c), where approximately 200 SNPs were involved in polygenic stabilizing selection (Fig. S4a). Regarding the causal SNPs, SNPs responsible for positive FDS showed strong population structures (*G*_st_ *>* 0.8) with low heterozygosity (*H*_t_ *<* 0.05; Fig. S4b) because positive FDS disrupts polymorphisms within a population. In contrast, since negative FDS maintains polymorphisms within a population, SNPs responsible for negative FDS had weak population structures (*G*_st_ *<* 0.2) with high heterozygosity (*H*_t_ *>* 0.35; Fig. S4b).

We implemented the single-locus model Equation (2) as a linear mixed model (LMM) for GWAS (Appendix S2) and evaluated the performance of LMMs in terms of causal polymorphism detection and effect size estimates (Fig. 4). The power to detect negative and positive FDS was strong (median AUC *>* 0.8) in both split and continuous settings (Fig. 4a, b, d and e). The direction of the FDS matches the estimated sign of *β*_2_ (Fig. 4c and f). In contrast, our method had almost no power to detect overdominance (Fig. 4b and e). Compared with overdominance, spatiotemporally varying selection was more likely, but the power remained weak (median AUC *<* 0.6) and its estimated coefficients had a median value of almost zero (Fig. 4b-c and e-f). These results indicate that our method retains the power to detect negative and positive FDS in GWAS, with other types of balancing selection less likely confounded.

**Figure 4:**
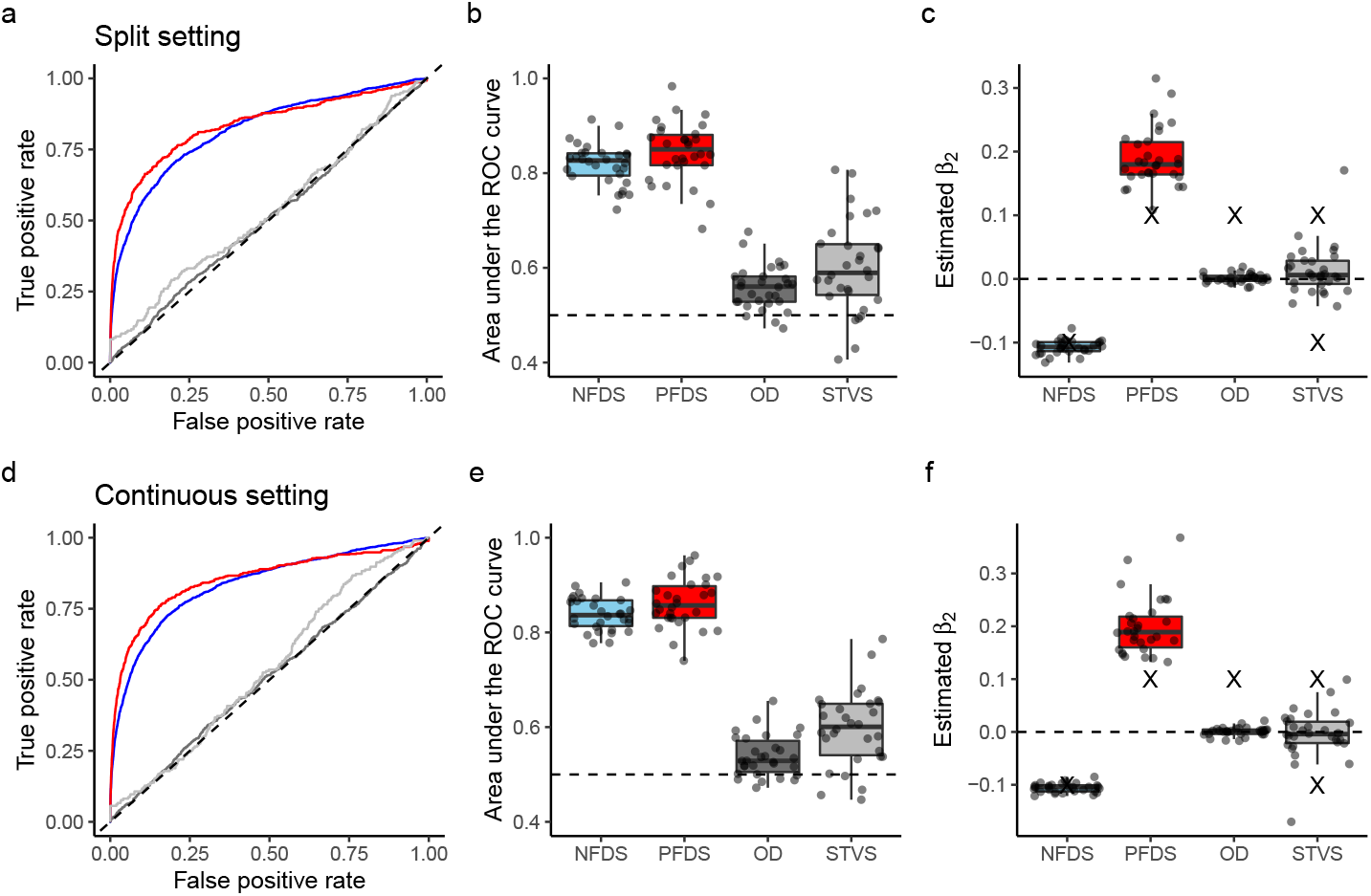
Performance of linear mixed models to estimate four types of simulated selection: NFDS, negative frequency-dependent selection (blue); PFDS, positive frequency-dependent selection (red); OD, overdominance (dark gray); and STVS, spatiotemporally varying selection (light gray). The top and bottom panels show the results of the split and continuous settings, respectively (Fig. 1a). The left panels (a) and (b) show the receiver operating characteristic (ROC) curve, which indicates the relationship between the true positive rate and false positive rate. The middle panels (b) and (e) show the area under the ROC curve (AUC). Dashed lines at AUC = 0.5 indicate no power to detect causal single nucleotide polymorphisms (SNPs). The right panels (c) and (f) show the estimated *β*_2_ of causal SNPs, where negative and positive values indicate negative and positive FDS, respectively. Cross marks indicate the true simulated magnitude of *β*_2_. Boxplots show the median by a center line, upper and lower quartiles by box limits, and 1.5× interquartile range by whiskers.

The performance of LMMs was compared with that of standard LMs (Fig. 4, Fig. S5). In terms of AUC, LMMs outperformed LMs in the detection of negative FDS (Fig. 4b and e, Fig. S5b and e). Although LMs and LMMs exhibited similar performances for positive FDS, LMs overestimated the true strength of positive FDS more than LMMs (Fig. 4c and f, Fig. S5c and f). LMMs retained a slight power to detect polygenic stabilizing selection (weakly concave ROC curve; Fig. S6a and d), whereas LMs had almost no power to detect the stabilizing selection (almost flat ROC curve; Fig. S7a and d). These results suggest that LMMs are better suited to the proposed method because they can more efficiently capture negative FDS or prevent overestimation of the strength of positive FDS.

### 3.4 Field GWAS of the branch number in *A. thaliana*

To examine feasibility using a real GWAS dataset, we finally applied an LMM for field GWAS of the branch number in *A. thaliana* (Fig. 5) under the continuous setting (Fig. 1a lower). QQ-plots exhibited little inflation of the observed *p*-value score against the expected score (Fig. S8). Of 561 plants, 181 bolted 2 weeks after they were transferred to the field in July. The proportion of branch number variation explained by self-genotypes 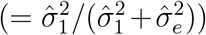 was 0.68, indicating high heritability of the fitness component. For the self-genotype effects *β*_1_ (i.e., standard GWAS), we detected no significant SNPs above the Bonferroni threshold but found a peak on the top of chromosome 4 (Fig. 5a; Table S4a). The top of chromosome 4 is known to encompass a flowering QTL and natural variation in the *FRI* (Aranzana *et al*., 2005). GO enrichment analysis detected no significant annotations at a false discovery rate of *<* 0.05.

**Figure 5:**
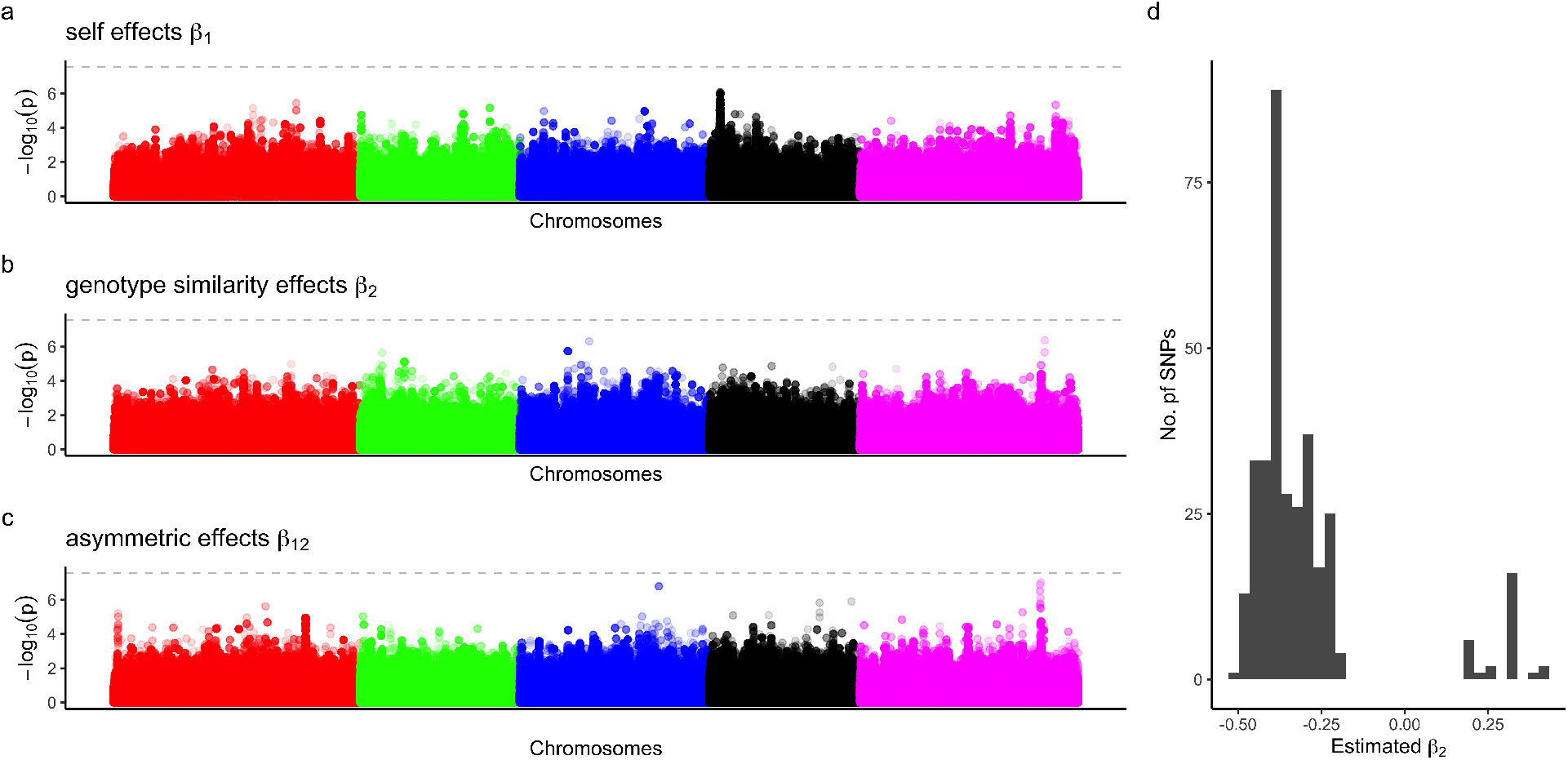
Genome-wide association studies of branch number in field-grown *Arabidopsis thaliana* under the continuous population setting (Fig. 1a lower). The results of linear mixed models are shown. (a, b, and c) Manhattan plots for self-genotype effects, genotype similarity effects, and asymmetric effects, respectively. Horizontal dashed lines indicate *p*-value = 0.05, after Bonferroni correction. (d) Histogram of estimated *β*_2_ among SNPs exhibiting *p*-values *<* 0.0001. Negative and positive *β*_2_ infer loci responsible for negative and positive FDS, respectively.

For the genotype similarity effects *β*_2_, we also found no significant SNPs but top-scoring SNPs on chromosomes 3 and 5 (Fig. 5b). Within 10 kbp near the top-scoring SNP on chromosome 3, we observed the *GAE6* gene, which encodes a UDP-D-glucuronate 4-epimerase involved in pectin biosynthesis, cell wall integrity, and immunity to pathogens (Bethke *et al*., 2016). To elucidate the genome-wide patterns of *β*_2_ and candidate genes, we focused on SNPs exhibiting *p*-values of *<* 10^−4^, corresponding to <0.025 percentiles. Of the 254 SNPs selected, 195 and 59 showed negative and positive *β*_2_, respectively (Fig. 5d). Genes involved in plant immunity and resistance, such as *GAE6, ARGONAUTE 4* (*AGO4*), and *ACTIVATED DISEASE RESISTANCE 1* (*ADR1*), were observed within 10 kb near the SNPs showing negative *β*_2_ at *p*-value *<* 0.0001 (Table S4b). In contrast, *AVRRPT2-INDUCED GENE 1* (*AIG1*) was only a resistance-related gene observed near the SNPs showing positive 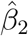 at *p*-values of *<* 0.0001 (Table S4b). GO enrichment analysis found no significant GO annotations at a false discovery rate of *<* 0.05.

We also tested the asymmetric effects *β*_12_ for all SNPs by detecting a QTL on chromosome 5 near the Bonferroni threshold (Fig. 5c). These SNPs exhibited positive *β*_2_ and *β*_12_ (i.e., 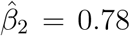 and 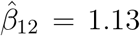 on chromosome 5 at position 22382673: Table S4c), indicating positive effects of a reference allele on absolute fitness together with positive FDS on relative fitness. Genes potentially related to growth were located near this top-scoring SNP, including *SUMO2* and *SUMO3*. GO enrichment analysis found no significant GO annotations at a false discovery rate of *<* 0.05.

To compare the LMMs and LMs, we applied standard linear models for the branch number (Fig. S9). For the self-genotype effects, a large number of SNPs showed larger -log_10_(*p*-values) scores than the Bonferroni threshold (Fig. S9a). The observed *p*-values for the self-genotype effects were greater than expected (Fig. S10a), showing that LMs should not be used for GWAS of the branch number. We also observed a slight inflation in the observed *p*-values for the genotype similarity and asymmetric effects (Fig. S10b-c). Consistent with the results of LMMs, the estimate sign of 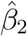 indicated that negative FDS rather than positive FDS was more likely observed among the top-scoring SNPs (Fig. S9d). Combined with the simulation above, these empirical results suggest that LMMs are more suitable for GWAS than LMs. The line of GWAS simulation and application shows that our method can also be used to screen polymorphisms associated with FDS (see also the discussion “Applicability for GWAS”)

## 4 Discussion

### 4.1 Selection gradient along genotype similarity

Selection gradient analysis is a powerful approach for empirical studies to quantify selection in action (Lande and Arnold, 1983; Mitchell-Olds and Shaw, 1987; Chong *et al*., 2018). By incorporating genotype similarity as a pseudo-trait, we propose a linear regression that simplifies a pairwise interaction model of FDS. The single-locus analysis of *A. halleri* and *I. elegans* suggests that our method is applicable to plants and animals in natural or semi-natural fields. As the covariate of genotype similarity 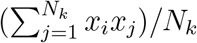 denotes how similar (or dissimilar) the neighbor compositions are with the focal individual, conclusions are expected to be the same as we regress fitness components on the frequency of other morphs. Empirical studies have often tested the effects of morph frequency on fitness in the subset data of each morph (McCauley and Brock, 1998; Bennington and Stratton, 1998; Sato and Kudoh, 2017) or evaluated relative fitness without monomorphic subpopulations (Gigord *et al*., 2001; Takahashi *et al*., 2010). In contrast to the analysis of partial data, the proposed method deals with a full dataset for statistical tests of the coefficient *β*_2_, which determines the direction and strength of the symmetric FDS.

Even when FDS is asymmetric between two alleles, another coefficient *β*_12_ helps us infer an accurate form of FDS on absolute fitness. Although the multiplicative model [Equation (3)] requires the additional estimation of *β*_12_, we could still analyze the full dataset without using the subset data of each morph. Practically, we should first test *β*_12_ using the multiplicative model and then test *β*_2_ using the linear model [Equation (2)] if *β*_12_ is not significant. The main effects 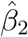 infer negative or positive FDS on relative fitness, whereas the coefficient of the asymmetric effect 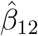 modulates the fitness slope along the allele frequency (Fig. 2). Despite the increased complexity due to the interaction term *β*_12_, the direction of FDS on the relative fitness can be simply interpreted by estimation of the main effect *β*_2_.

Alteration of the population-level mean fitness is another remarkable outcome of the pairwise interaction model (Cockerham *et al*., 1972; Asmussen and Basnayake, 1990; Schneider, 2008). Based on the simplified model of pairwise interactions, our method provides an additional inference of the relationships between the population-level mean fitness and allele frequency (Fig. 2). For instance, with symmetric negative FDS, the mean fitness is expected to increase more in polymorphic than in monomorphic populations (Fig. 2a). The line of empirical studies on *I. elegans* has reported such an increased population-level mean fitness at an intermediate frequency (Takahashi *et al*., 2014) as well as negative FDS on female color polymorphisms (Le Rouzic *et al*., 2015). The results of our reanalysis show symmetric negative FDS (i.e., 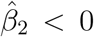; Table 1b) and a consequent increase in population-level mean fitness at an intermediate frequency in *I. elegans* (Fig. 3b). Although the reanalysis of *A. halleri* data shows the joint action of directional selection and biased frequency on its equilibrium, negative FDS still maintained the dimorphism and very slightly increased population-level mean fitness at an intermediate equilibrium frequency (Fig. 3a). Combined with the previous evidence, the present method provides an empirical approach to understand how FDS increases population-level mean fitness.

### 4.2 Applicability for GWAS

By incorporating the population structure as random effects, we extended our method to LMMs that have often been used in GWAS (Kang *et al*., 2008) (see also Appendix S2). Our simulations suggest that LMMs improve the power to detect causal polymorphisms or prevent us from exaggerating effect-size estimates of FDS. However, caution should be exercized regarding the genetic structure of the loci underlying positive or negative FDS. As positive FDS disrupts polymorphisms, their selected loci likely showed low heterozygosity and strong population differentiation (Fig. S4). While LMMs could deal with the population structure, their effect-size estimates were still larger than the true signals (Fig. 4c and f). This might be due to a similar genetic structure between the neutral and selected loci when a genome underwent positive FDS. In contrast, negative FDS results in high heterozygosity by maintaining polymorphisms within a population (Fig. S4), where LMMs performed better than LMs by separating the neutral population structure and loci subject to negative FDS. Furthermore, the signals of negative and positive FDS were well distinguished from those of overdominance and spatiotemporally varying selection.

The application to *A. thaliana* accessions showed a genome-wide excess of negative FDS and a few loci underlying asymmetric positive FDS. This result seems plausible because polymorphic loci are more likely to persist under negative FDS than positive FDS. In the present study, we found candidate genes that might be involved in conferring plant resistance to pathogens. Several studies have reported negative effects of FDS on plant resistance to pathogens (Antonovics and Ellstrand, 1984; Brunet and Mundt, 2000). In contrast, such resistance genes were not found near the loci responsible for asymmetric positive FDS. We also found growth-related candidate genes near the loci associated with asymmetric positive FDS. Plant competition is known to exert asymmetric and positive frequency-dependent effects from tall to short plants (Weiner, 1990), where growth-related loci are more likely to be observed than defense-related loci. However, the top-scoring SNPs were still below the genome-wide threshold of significance. Thus, experimental studies using single-gene mutants are necessary to validate FDS on genes related to plant resistance or competition.

### 4.3 Potential limitation

Pairwise interaction models have been extensively analyzed in relation to one locus with multi-alleles (Schneider, 2006; Trotter and Spencer, 2007) and multi-loci with multiple alleles (Schneider, 2010), in addition to the model of one locus with two alleles (Cockerham *et al*., 1972; Asmussen and Basnayake, 1990; Schneider, 2008). At the expense of wide applicability, our method was too simplified to reflect all the theoretical features of pairwise interaction models. For example, the combination of overdominance and FDS at the same locus or epistasis among loci responsible for FDS was not considered in the current regression model. Even the one-locus two-allele model can make fitness functions have multiple equilibria when it involves the additive action of FDS on a quantitative trait (Appendix S3), yet more complex outcomes may arise from the realistic genetic architecture. In nature, FDS may act on more than two co-dominant alleles at a single locus, such as the S-allele system in plant self-incompatibility (Hatakeyama *et al*., 1998; Shimizu and Tsuchimatsu, 2015). Multi-state extension of the Ising model, which is known as the Potts model (Potts, 1952), may deal with this situation if plausible mechanisms of biological inheritance can be incorporated.

### 4.4 Conclusions

The present study offers an effective way to resolve fitness-genotype associations with respect to FDS. Our phenotype-driven approach can distinguish between positive and negative FDS based on direct observations of fitness. In molecular population genetics, phenotype-free methods are available to detect signatures of past balancing selection (Siewert and Voight, 2017). While fitness measurements are labor-intensive, selection gradient analysis has an advantage in quantifying ongoing FDS. Now that genome information is accumulating in wild organisms (Lewin *et al*., 2018), organismal biologists may utilize genome data for a deeper understanding of the maintenance of polymorphism. A joint approach using genomics and fitness evaluation would enable future studies to unveil FDS throughout the genome.

## Acknowledgments

The authors thank M. Yamazaki, A. Morishima, and all members of the Shimizu group for their assistance with the setup of the *A. thaliana* experiment; H. Kudoh for discussions during the *A. halleri* study; and J. Bascompte and S.E. Wuest for comments on the draft. Y.S. thanks A.J. Nagano for hosting him as a guest researcher at Ryukoku University. This study was supported by the Japan Society for the Promotion of Science (JSPS) KAKENHI (Grant No. 20K15880) and Japan Science and Technology Agency (JST) PRESTO (JPMJPR17Q4) grants (Y.S.) and Swiss National Science Foundation (31003A_182318) and JST CREST (JPMJCR16O3) grants (K.K.S.). The fieldwork at Zurich was supported by the University of Zurich via the University Research Priority Program for “Global Change and Biodiversity.” The authors declare no conflict of interest.

## Author Contributions

Y.S. developed the model, analyzed the data, and wrote a draft. Y.T. provided the damselfly data, discussed the results, and contributed to the conceptualization regarding FDS. C.X. and Y.S. conducted the field experiment using *A. thaliana*. Y.S. and K.K.S. designed the study, and revised the manuscript with inputs from the co-authors.

## Data Availability Statement

The source codes and original data generated by this study are available in the GitHub repository (https://github.com/yassato/RegressionFDS).

## Supplementary Materials

### Appendix S1. Metropolis algorithm with Mendelian inheritance

The forward problem of the Ising model is to optimize the individual fitness *w*_*i*_ with the given selection coefficients *s*_1_ and *s*_2_. Figure 1 shows two contrasting cases, where one represents short-range interactions in a lattice space while the other shows uniform interactions within a series of split populations. Individual fitness is defined by Equation (1) in the main text as 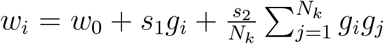. The magnetic interaction was defined in the Ising model as *H* = −*J* Σ_<*i,j*>_*g*_*i*_*g*_*j*_ + *η* Σ*g*_*i*_, where the total energy *H* can be regarded as the population sum of fitness Σ*w*_*i*_, magnetic interaction coefficient *J* as the total interaction strength *s*_2_*N*_*k*_, and external magnetic force *η* as the total strength of directional selection *s*_1_*N*_*k*_.

To represent inheritance and selection, we updated the genotype *g*_*i*_(*j*) and fitness *w*_*i*_ based on the Metropolis algorithm and Mendelian inheritance. The individual fitness *w*_*i*_ was updated following the Metropolis algorithm (Metropolis *et al*., 1953), which has often been used in a series of stochastic sampling methods, such as the Markov chain Monte Carlo method (Bishop, 2006). Here, we consider a change in the genotype *g*_*i*_ from generation *t* to *t* + 1. Let 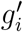be a proposed genotype for *t* + 1, and let 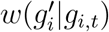 be a conditional fitness with a given genotype *g*_*i,t*_ at *t*. Then, 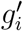 can be accepted/rejected based on its likelihood ratio on the current fitness w(*g*_*i,t*_) as

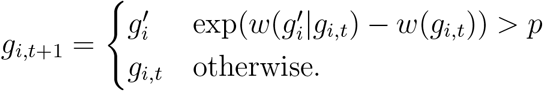

where scalar *p* is sampled from a uniform distribution as *p* ∼ Unif(0, 1). This update process may mimic selection to some extent of stochasticity.

To incorporate Mendelian inheritance into the forward problem of the Ising model, the transition from *g*_*i,t*_ to *g*_*i,t*+1_ was weighted by genotype segregation among the offspring. When two genotypes *g*_*i,t*_ and *g*_*j,t*_ crossed each other at generation *t*, we could expect nine possible combinations among parental genotypes, as shown in Table S1.

**Table S1:** Cross tables among AA, Aa, and aa genotypes.

| $g_i / g_j$ | AA | | A | | aa | |
| --- | --- | --- | --- | --- | --- | --- |
| AA | AA | AA | AA | Aa | Aa | Aa |
|  | AA | AA | AA | Aa | Aa | Aa |
| Aa | AA | AA | AA | Aa | Aa | Aa |
|  | Aa | Aa | Aa | aa | aa | aa |
| aa | Aa | Aa | Aa | aa | aa | aa |
|  | Aa | Aa | Aa | aa | aa | aa |

The probability of sampling AA, Aa, and aa at *t* + 1 generation is denoted as *P*_AA,*t*+1_, *P*_Aa,*t*+1_, and *P*_aa,*t*+1_, respectively. Let *f*_AA_, *f*_Aa_, and *f*_aa_ be the frequencies of genotypes AA, Aa, and aa within a population with *f*_AA_, *f*_Aa_, and *f*_aa_. In summary, the transition from generation *t* to *t* + 1 is expressed as

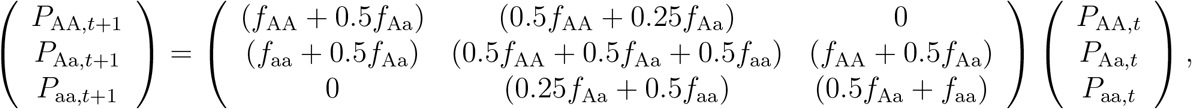

where the elements of the transition matrix were calculated from the three genotype frequencies and the segregation ratio of Mendelian inheritance. The zero elements pose a constraint where one homozygote cannot turn into another homozygote in a single generation. Given that the sum of the three genotype frequencies was 1 as *f*_AA_ +*f*_Aa_ +*f*_aa_ = 1, the probability of remaining as a heterozygote was 0.5. Therefore, the outcome from the modified Metropolis algorithm was expected to be qualitatively the same as a random proposal of three genotypes, but quantitatively different in the excess of heterozygotes within a population.

**Figure S1:**
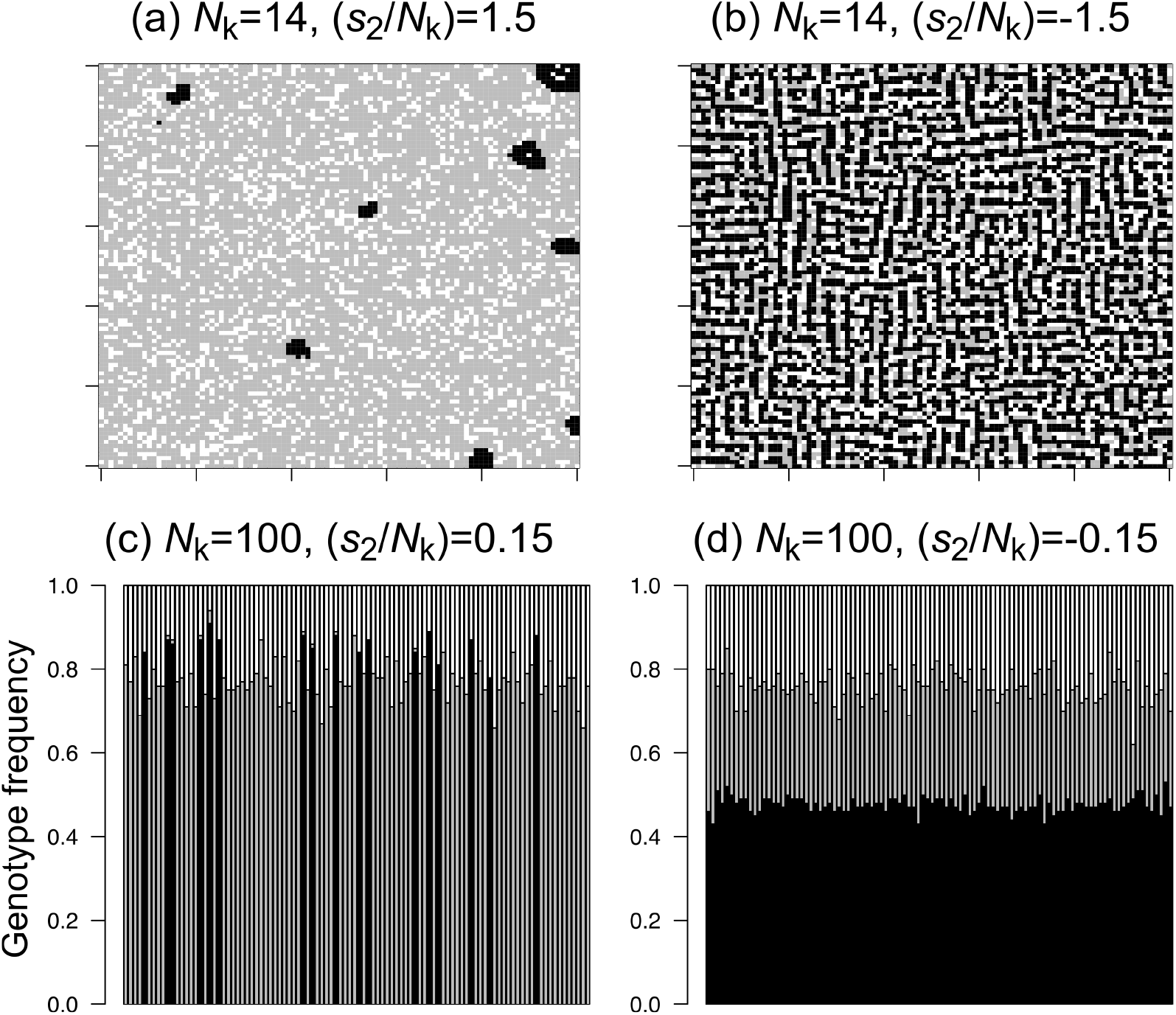
Numerical simulations maximizing the fitness 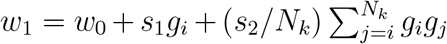 (Appendix S1) for 100 generations. White, gray, and black indicate the AA, Aa, and aa genotypes, respectively. Top panels (a, b) represent plant genotype distributions across a 100 × 100 continuous lattice space when interactions are restricted to the second nearest neighbors (*N*_*k*_ = 14). Each grid corresponds to an individual. Bottom panels (c, d) represent the genotype frequencies among 100 split populations composed of 100 individuals each. Each vertical bar corresponds to a population. Left panels (a, c) simulate positive frequencydependent selection (FDS), while right panels (b, d) simulate negative FDS. The base fitness and directional selection coefficient were set at *w*_0_ = 0 and *s*_1_ = 10^−4^ for all simulations, while the subpopulation size *N*_*k*_ and interaction strength per individual *s*_2_*/N*_*k*_ were changed.

Figure S1 shows the results of the numerical simulations with the three genotypes encoded as *g*_*i*(*j*)_ ∈ {AA, Aa, aa} = {+1, +1, -1}. The three genotypes were well mixed and maintained in a continuous space when *s*_2_ *<* 0 (Fig. S1b), whereas several clusters of the aa genotype were observed when *s*_2_ *>* 0 (Fig. S1a). Three genotypes were also maintained at an intermediate frequency in a split space when *s*_2_ *<* 0 (Fig. S1d), whereas the allele frequency was heavily biased when *s*_2_ *>* 0 (Fig. S1c). The numerical simulations show that the sign of *s*_2_ likely corresponded to the direction of the frequency-dependent selection (FDS) in a continuous and split space.

### Appendix S2. Mixed model extension

To implement GWAS, we modified Equations (2) and (3) as a linear mixed model (LMM) that considered genetic relatedness as a random effect (Kang *et al*., 2008). In terms of Henderson’s mixed model (Henderson *et al*., 1959), such GWAS models have the same structure as phylogenetic comparative methods that analyze interspecific phenotypic variation among phylogenetic trees (Kang *et al*., 2008; Hadfield and Nakagawa, 2010). According to Hadfield and Nakagawa (2010), we designated the genetic-related matrix as **A** and introduced random effects *u*_*i*_ to Equation (2) as follows:

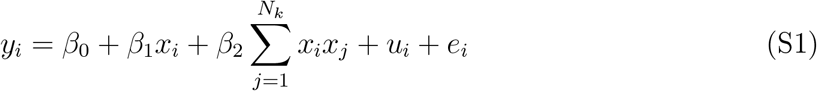

where a vector including *u*_*i*_ for *n* individuals followed a normal distribution as *u*_*i*_ ∈ **u** and 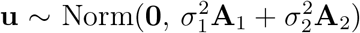. The residual *e*_*i*_ is expressed as *e*_*i*_ ∈ **e** and 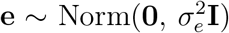. The *n* × *n* variance–covariance matrices denote the self-genetic relatedness or the entire genotype similarity among *n* individuals as 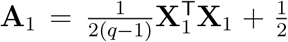and 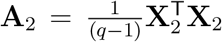, where *q* denotes the number of loci. The elements of *n* individuals × *q* loci matrix **X**_1_ consist of explanatory variables of the self-genotype values. As we defined *x*_*i*(*j*)_ = (−1, 1), the genetic-related matrix **A**_1_ was scaled to represent the proportion of loci shared among *n* × *n* individuals. The elements of *n* individuals × *q* loci matrix **X**_2_ consist of the genotype similarity as

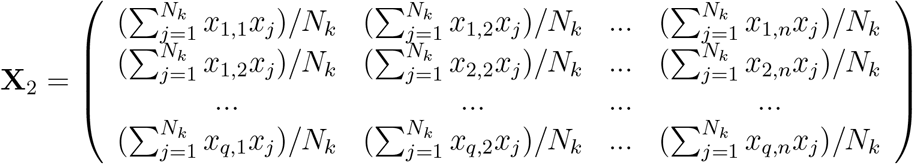

where the *n* × *n* matrix **A**_2_ indicates a sample structure related to genotype similarity. The variance component parameters 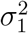 and 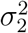 determine the relative contributions of **A**_1_ and **A**_2_ to the vector of random effects **u**.

To incorporate asymmetric FDS into GWAS, we considered a sample structure based on the asymmetric FDS in LMM. Here, we extended Equation (3) into LMM as

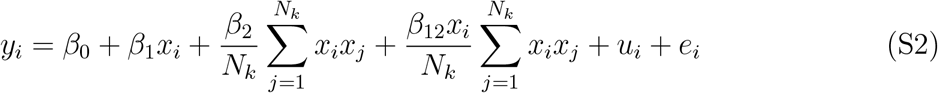

where the random effect *u*_*i*_ is redefined as *u*_*i*_ ∈ **u** and 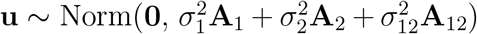. The additional *n* × *n* variance-covariance matrix **A**_12_ denotes a sample structure because of the asymmetric effects among *n* individuals as 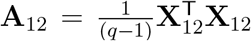. The elements of *n* individuals × *q* loci matrix **X**_12_ consist of explanatory variables for the asymmetric effects as follows:

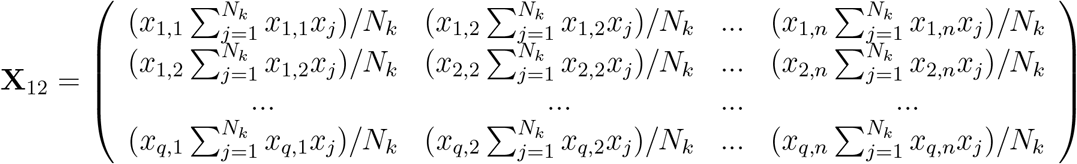

The additional parameter of the variance component 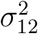compares the relative importance of the asymmetric effects with those of the self-genotype effects 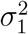 and genotype similarity effects 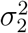. Additionally, the individual-level formula Equation (S1) can also be converted into a common matrix form (Henderson *et al*., 1959) as follows:

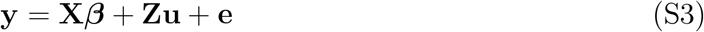

where **y** is *n* × 1 fitness vector with *y*_*i*_ ∈ **y**; **X** is a matrix of fixed effects, including a unit vector, self-genotype *x*_*i*_, genotype similarity covariate 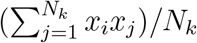, and other covariates for *n* individuals; ***β*** is a vector that included coefficients of the fixed effects; **Z** is a design matrix allocating individuals to a genotype; **u** is the random effect as 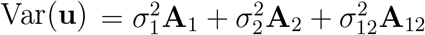; and **e** is the residual as 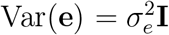.

To efficiently solve LMMs, we first estimated 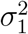 and 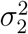 without any fixed effects (i.e., null model) using the average-information restricted maximum likelihood method implemented

in the gaston package (Perdry and Dandine-Roulland, 2020). Then, to compare the significance of *β*_1_ and *β*_2_ to the null model, we solved Equation (S3) using fast approximation by eigenvalue decomposition on a random effect matrix 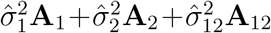. The likelihood ratio test was used to compare the models with and without *β*_2_. The standard GWAS is a subset of Equation (S1) when *β*_2_ = 0 and 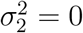 (Sato *et al*., 2021b); thus, we set *β*_2_ and 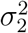 to zero when testing *β*_1_. Such stepwise likelihood ratio tests were essential for conservative tests of each parameter in Equations (S1) and (S2) because the self-genotype variable *x*_*i*_ and the genotype similarity variable 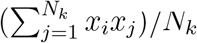 were strongly correlated when the MAF was very small (Sato *et al*., 2021b). Stepwise likelihood ratio tests were implemented using the rNeighborGWAS package version 1.2.4 (Sato *et al*., 2021b). The nei_lmm() function was applied for power analysis and GWAS in the main text. The “asym = TRUE” option was chosen when testing the asymmetric effects *β*_12_.

### Appendix S3. Fitness function under symmetric and asymmetric FDSs

To analyze the regression model Equation (3) as a fitness function of allele frequency, we considered a single diallelic locus in an ideal population where (i) diploid individuals were randomly mating, (ii) uniformly interacting, and (iii) the population size was sufficiently large (i.e., *N* → ∞). We also assumed no maternal or paternal effects on fitness such that the genotypes Aa and aA could not be distinguished. To concentrate on the mean trends of the model, we neglected the residuals as *e*_*i*_ = 0. Replacing *N*_*k*_ into *N*, we redefined Equation (3) as follows:

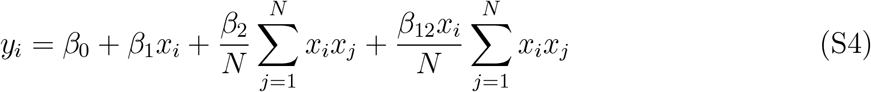

where the trait value *y*_*i*_ corresponds to the fitness value *w*_*i*_ for individual *i*, the intercept *β*_0_ corresponds to the base fitness *w*_0_, the self-genotype coefficient *β*_1_ corresponds to the directional selection coefficient *s*_1_, and the coefficients *β*_2_ and *β*_12_ correspond to the selection coefficients related to FDS. The second term can also be transformed for simplicity as 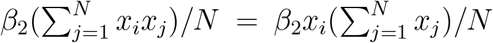, and the third term, 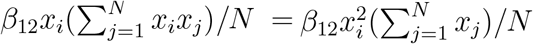. Provided 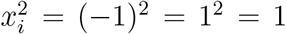, the third term represents a selection gradient due to the relative abundance of one allele in a neighborhood, irrespective of the self-genotype of a focal individual *i*.

We then transformed Equation (S4) into a function of the frequency of A alleles *f*, where the frequency of an allele was defined conversely as 1 − *f*. When all the individuals were randomly interacting in a panmictic population, the interaction strength between *i* and *j* depended on the frequencies of AA, Aa, or aa genotypes derived from allele frequency. Thus, the fitness values of the three genotypes, *y*_AA_(*f*), *y*_Aa_(*f*), and *y*_aa_(*f*), are functions of allele frequency *f*. For convenience, we suppressed the dependence on *f*, unless necessary. The ratio of AA, Aa, and aa genotypes within the panmictic population was given by AA:Aa:aa = *f* ^2^ : 2*f* (1 − *f*): (1 − *f*)^2^. Assuming the aforementioned ideal population, we designated all combinations of interactions among AA, Aa, and aa genotypes and weighted the interactions based on their genotype frequencies. Table S2 lists the interaction strength weighted by the genotype frequency for symmetric and asymmetric effects. Based on these cross tables (Table S2), we redefined Equation (S4) for the three genotypes as:

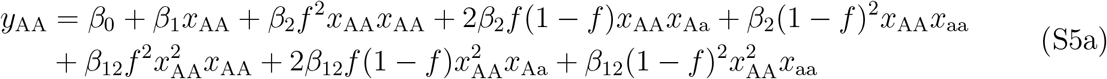

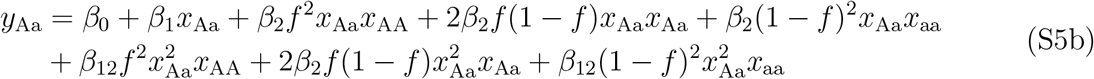

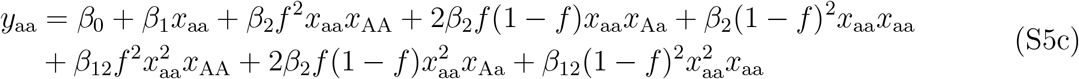

**Table S2:** Cross tables showing the strength of pairwise interactions between the focal genotype *x*_*i*_ and counterpart *x*_*j*_ in a randomly interacting and mating population.

(a) Symmetric effects
| $x_j \setminus x_i$ | $x_{AA}$ | $x_{Aa}$ | $x_{aa}$ |
| --- | --- | --- | --- |
| $x_{AA}$ | $\beta_2 f^2 x_{AA} x_{AA}$ | $\beta_2 f^2 x_{Aa} x_{AA}$ | $\beta_2 f^2 x_{aa} x_{AA}$ |
| $x_{Aa}$ | $2\beta_2 f(1-f) x_{AA} x_{Aa}$ | $2\beta_2 f(1-f) x_{Aa} x_{Aa}$ | $2\beta_2 f(1-f) x_{aa} x_{Aa}$ |
| $x_{aa}$ | $\beta_2 (1-f)^2 x_{AA} x_{aa}$ | $\beta_2 (1-f)^2 x_{Aa} x_{aa}$ | $\beta_2 (1-f)^2 x_{aa} x_{aa}$ |

(b) Asymmetric effects
| $x_j \setminus x_i$ | $x_{AA}$ | $x_{Aa}$ | $x_{aa}$ |
| --- | --- | --- | --- |
| $x_{AA}$ | $\beta_{12} f^2 x_{AA}^2 x_{AA}$ | $\beta_{12} f^2 x_{Aa}^2 x_{AA}$ | $\beta_{12} f^2 x_{aa}^2 x_{AA}$ |
| $x_{Aa}$ | $2\beta_{12} f(1-f) x_{AA}^2 x_{Aa}$ | $2\beta_{12} f(1-f) x_{Aa}^2 x_{Aa}$ | $2\beta_{12} f(1-f) x_{aa}^2 x_{Aa}$ |
| $x_{aa}$ | $\beta_{12} (1-f)^2 x_{AA}^2 x_{aa}$ | $\beta_{12} (1-f)^2 x_{Aa}^2 x_{aa}$ | $\beta_{12} (1-f)^2 x_{aa}^2 x_{aa}$ |

We further weighted the genotype-level fitness values by allele frequency. The allele-level marginal fitness for A or an allele is then given by:

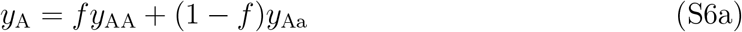

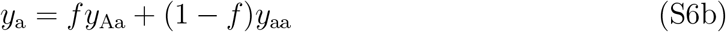

The population-level mean fitness was finally defined by the weighted mean of the marginal fitness as follows:

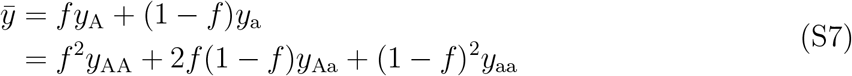

Consequently, we could analyze fitness functions by inputting specific values in the three genotype values *x*_AA_, *x*_Aa_, and *x*_aa_ throughout Equations (S5) and (S6).

Case 1. Complete dominance with random mating: Novel mutations are expected to be recessive during adaptive evolution. Empirical studies have reported FDS on dimorphic traits that often exhibit complete dominance of one over another allele (e.g., Takahashi *et al*., 2010; Sato and Kudoh, 2017; Goldberg *et al*., 2020). First, we considered the case in which the A allele was completely dominant over an allele, as encoded by *x*_*i*(*j*)_ ∈ {AA, Aa, aa} = {+1, +1, -1}. Here, we neglected the directional selection as *β*_1_ = 0 in Equation (S5) to focus on the fitness functions under FDS alone. Replacing the genotypes (*x*_AA_, *x*_Aa_, and *x*_aa_) according to their genotype values in Equation (S5) gave the fitness value to the three genotypes as

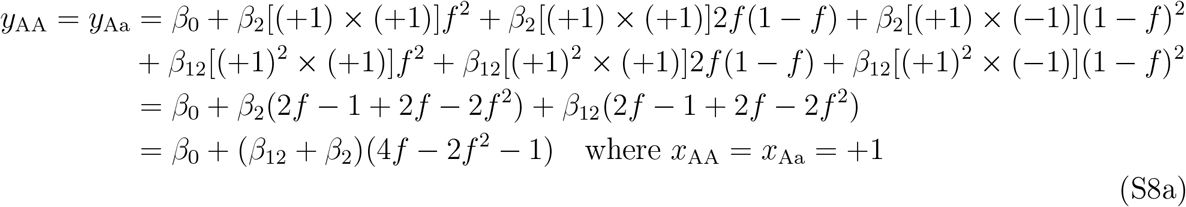

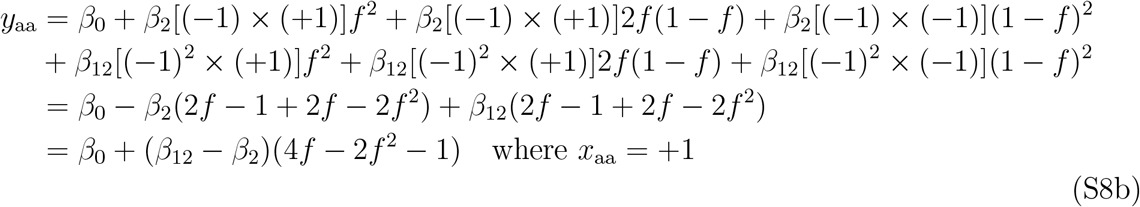

The marginal fitness following Equations (S6a) and (S6b) was given by

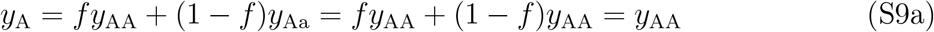

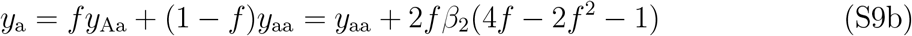

The relative fitness between A and a alleles was calculated as: *y*_A_−*y*_a_ = *y*_AA_−*y*_aa_−2*fβ*_2_(4*f* − 2*f* ^2^ − 1) = 2*β*_2_(1 − *f*)(4*f* − 2*f* ^2^ − 1). Solving *y*_A_ − *y*_a_ = 0 with respect to *f* within the range of (0,1) resulted in 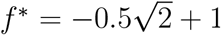, which showed a single stable or unstable state within *f* = (0, 1) in the case of complete dominance under FDS. The population-level mean fitness is finally given by

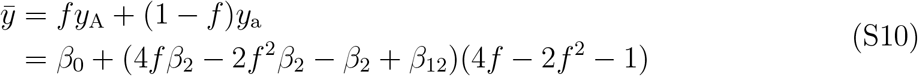

Figure 2 in the main text shows the marginal fitness [Equations (S9a) and (S9b)] and mean fitness [Equation (S10)]. The mean fitness was maximized at a stable equilibrium at the intermediate allele frequency under symmetric negative FDS, whereas it was minimized at an unstable equilibrium under symmetric positive FDS. When asymmetric FDS and complete dominance are involved, equilibria do not always match the maxima or minima of the mean fitness because of the nonlinearity of the marginal fitness in response to *f*. However, the mean fitness at the stable or unstable point was still higher or lower than expected compared with the weighted mean of the two monomorphic populations. A similar notion was suggested by pairwise interaction models in population genetics (Cockerham *et al*., 1972; Schneider, 2008).

Case 2. Additive effects with random mating: Although few empirical studies have reported FDS on quantitative traits, this case is of theoretical interest in the pairwise interaction model (Schneider, 2008). Considering the fitness value as a quantitative trait, we then analyzed the additive effects of A and a alleles on *y*_*i*_ as encoded by *x*_*i*(*j*)_ ∈ {AA, Aa, aa} = {+1, 0, -1}. Replacing the genotypes (*x*_AA_, *x*_Aa_, and *x*_aa_) according to their genotype values in Equations (S5) gave the fitness value to the three genotypes as:

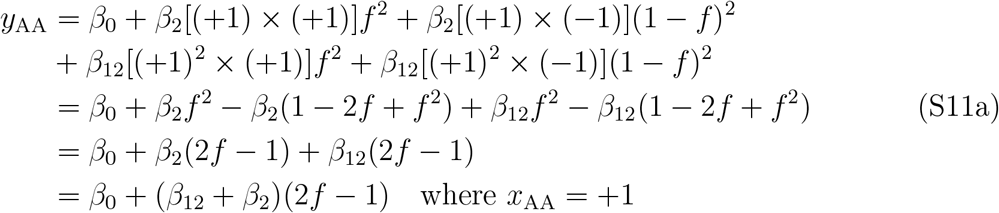

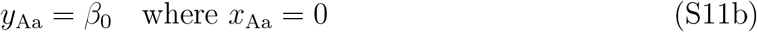

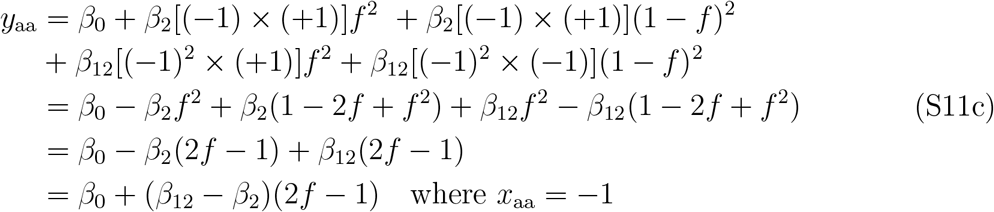

The fitness function for the two homozygotes AA and aa [i.e., Equations (S5a) and (S5c)] turned out to be linear in response to *f*. The marginal fitness following Equations (S6a) and (S6b) is then given by:

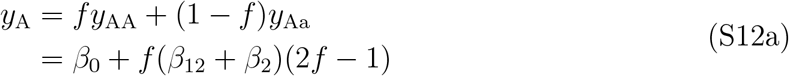

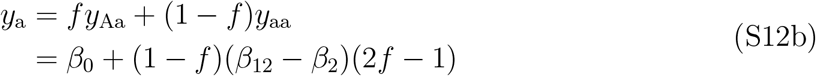

The relative fitness was calculated as *y*_A_ − *y*_a_ = (2*f* − 1)[*f* (*β*_12_ + *β*_2_) − (1 − *f*)(*β*_12_ − *β*_2_)] = (2*f* − 1)(2*β*_12_*f* − *β*_12_ + *β*_2_). Solving *y*_A_ − *y*_a_ = 0 with respect to *f* within a range of (0,1) yields *f* ^*^ = 0.5 and *f* ^*^ = 0.5(*β*_12_ − *β*_2_)*/β*_12_, showing that the additive action of FDS made multiple equilibria possible at the intermediate allele frequency *f* = (0, 1). The mean fitness following Equation (S7) is finally given by:

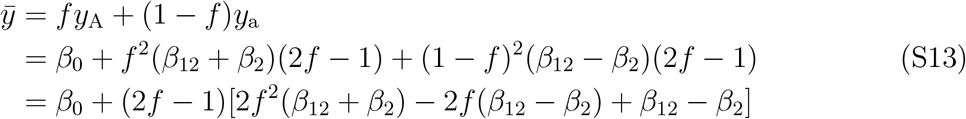

Figure S2 shows numerical examples of the marginal fitness [Equations (S12a) and (S12b)] and the mean fitness [Equation (S13)] in response to *f*. Similar to the case of complete dominance, the mean fitness was maximized or minimized at a stable or unstable equilibrium under the symmetric FDS (Fig. S2a and b). In contrast, a stable and unstable equilibrium occurred simultaneously under asymmetric FDS (Fig. S2c and f), where the maxima or minima of mean fitness did not always match the equilibria. This potential of multiple equilibria was also suggested by a one-locus two-allele model of pairwise interactions when it involved asymmetric FDS (Schneider, 2008).

Case 3. Asexual or inbred lines without mating: In common gardens or laboratory experiments, researchers arbitrarily distribute inbred accessions in space and retrieve individuals before mating (e.g., Schutz and Usanis, 1969; Sato *et al*., 2021b). This was also the case for the field GWAS in the main text. Furthermore, ecological studies often focus on FDS at the phenotype level with asexual reproduction assumed (e.g., Takahashi *et al*., 2018). In these cases, heterozygosity was negligible, and the two homozygotes were encoded as *x*_*i*(*j*)_ ∈ {AA, aa} = {+1, -1} (Sato *et al*., 2021b). Let *f*_AA_ and *f*_aa_ be the frequency of AA and aa genotypes within a population, where *f*_AA_ + *f*_aa_ = 1. The fitness function for the AA or aa genotype is given by:

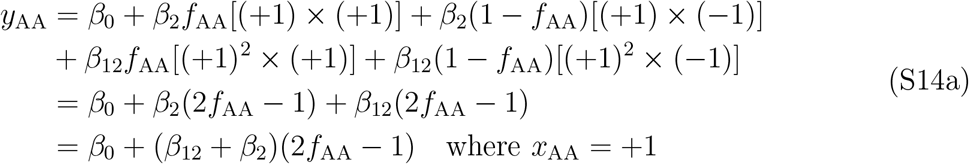

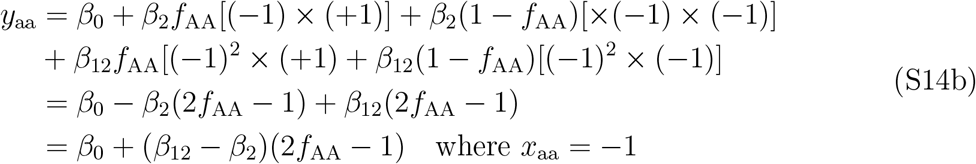

where the population-level mean fitness was given by its weighted mean as follows.

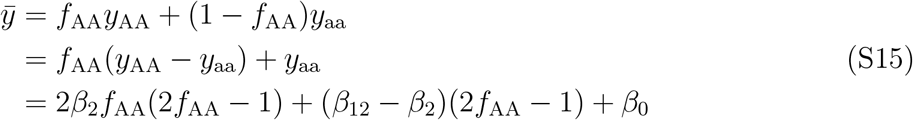

**Figure S2:**
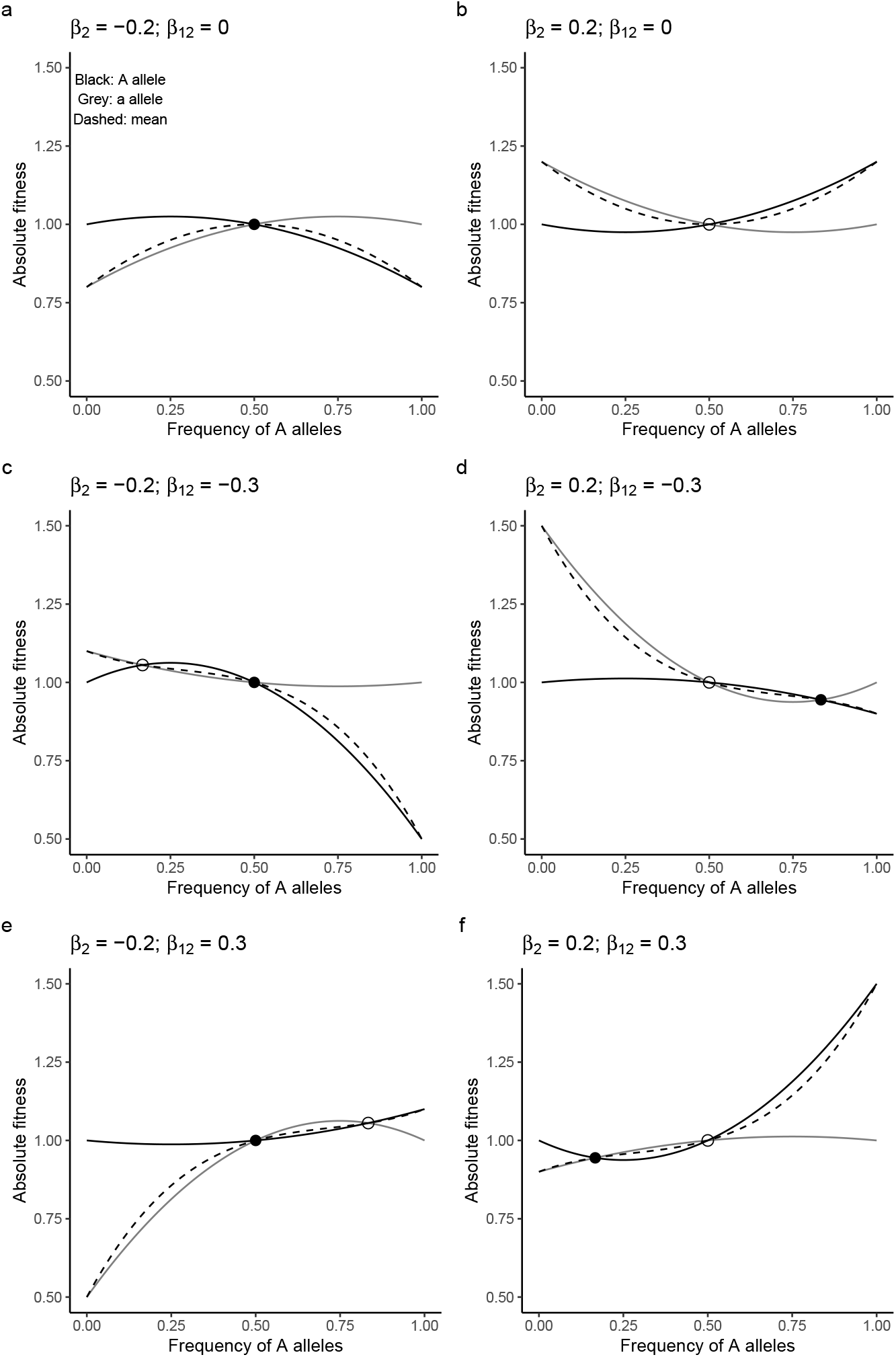
Numerical examples for the fitness values *y*_*i*_ in response to allele frequency when A and a alleles have additive effects on the fitness; that is, Equations (S12a) and (S12b) in Appendix S3. Black and gray lines indicate the marginal fitness of the A allele or an allele, respectively. The dashed curves indicate the mean fitness per population; that is, Equation (S13) in Appendix S3. (a) Symmetric negative frequency-dependent selection (FDS); (b) symmetric positive FDS; (c and e) asymmetric negative FDS; and (d and f) asymmetric positive FDS. Closed and open circles indicate a stable or unstable state, respectively. The base fitness and no directional selection were set at *β*_0_ = 1.0 and *β*_1_ = 0.0 for all panels.

Figure S3 shows the fitness function of the AA or aa genotype [Equations (S14a) and (S14b)] and population mean [Equation S15]. The fitness function of the two homozygotes was the same as that of the additive case described above. The mean fitness became simpler as the allele frequency corresponded to the genotype frequency. As this inbred case represented two genotypes with asexual reproduction, its conclusion was basically the same as that derived from game theoretical models (Takahashi *et al*., 2018).

In Figure 3, we present this inbred case with *β*_1_ ≠ 0, where the genotype fitness Equations (S14a) and (S14b) is rewritten as

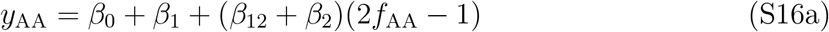

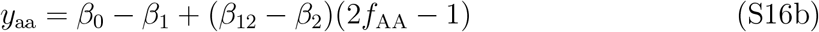

where the relative fitness is given by *y*_AA_ − *y*_aa_ = 2*β*_1_ + 2*β*_2_(2*f*_AA_ − 1). Solving *y*_AA_ − *y*_aa_ = 0 with respect to *f*_AA_ = (0, 1) yields 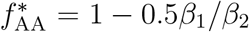. Therefore, the directional selection coefficient *β*_1_ may modify the equilibrium. When *β*_1_ ≠ 0, the mean fitness Equation (S15) can also be rewritten as:

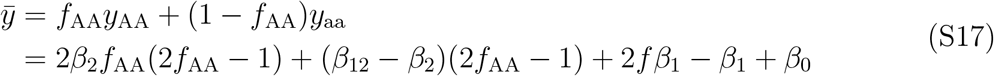

**Figure S3:**
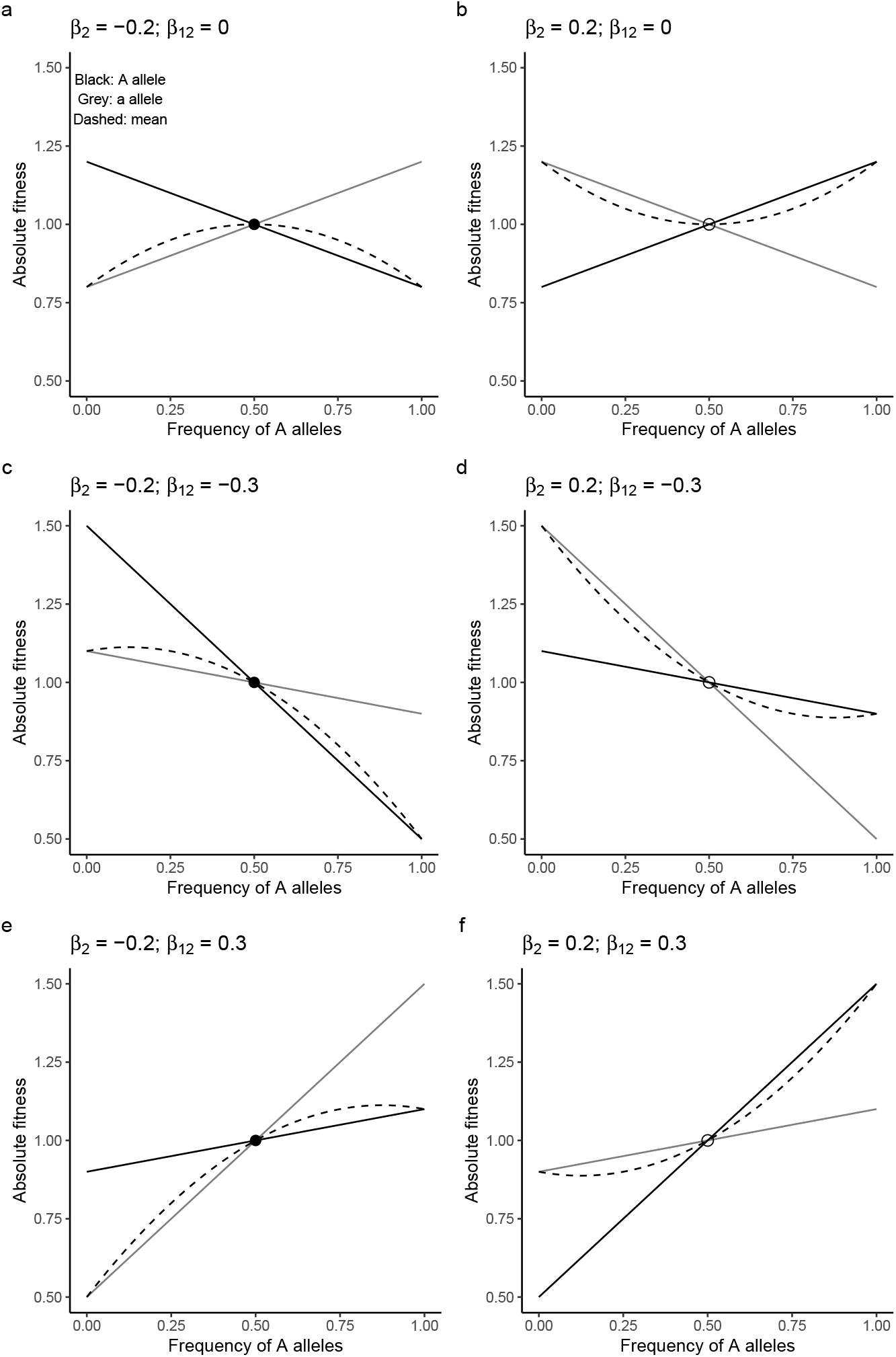
Numerical examples of fitness values *y*_*i*_ in response to allele frequency when only the AA and aa genotypes exist without mating [Equations (S14a) and (S14b) in Appendix S3]. Black and gray lines indicate the fitness functions for the AA and aa genotypes, respectively. The dashed curves indicate the mean fitness per population; that is, Equation (S15) in Appendix S3. (a) Symmetric negative frequency-dependent selection (FDS); (b) symmetric positive FDS; (c and e) asymmetric negative FDS; and (d and f) asymmetric positive FDS. Closed and open circles indicate a stable or unstable state, respectively. The base fitness and no directional selection were set at *β*_0_ = 1.0 and *β*_1_ = 0.0 for all panels.

### Supplementary Tables S3-S4 (see SuppTablesS3-S4.xlsx)

Table S3: List of plant accessions used for genome-wide association studies (GWAS) and their phenotypes.

Table S4: List of candidate genes related to self-genotype effects (a), genotype similarity effects (b), and asymmetric effects (c) on the branch number in *Arabidopsis thaliana*.

### Supplementary Figures S4–S10

**Figure S4:**
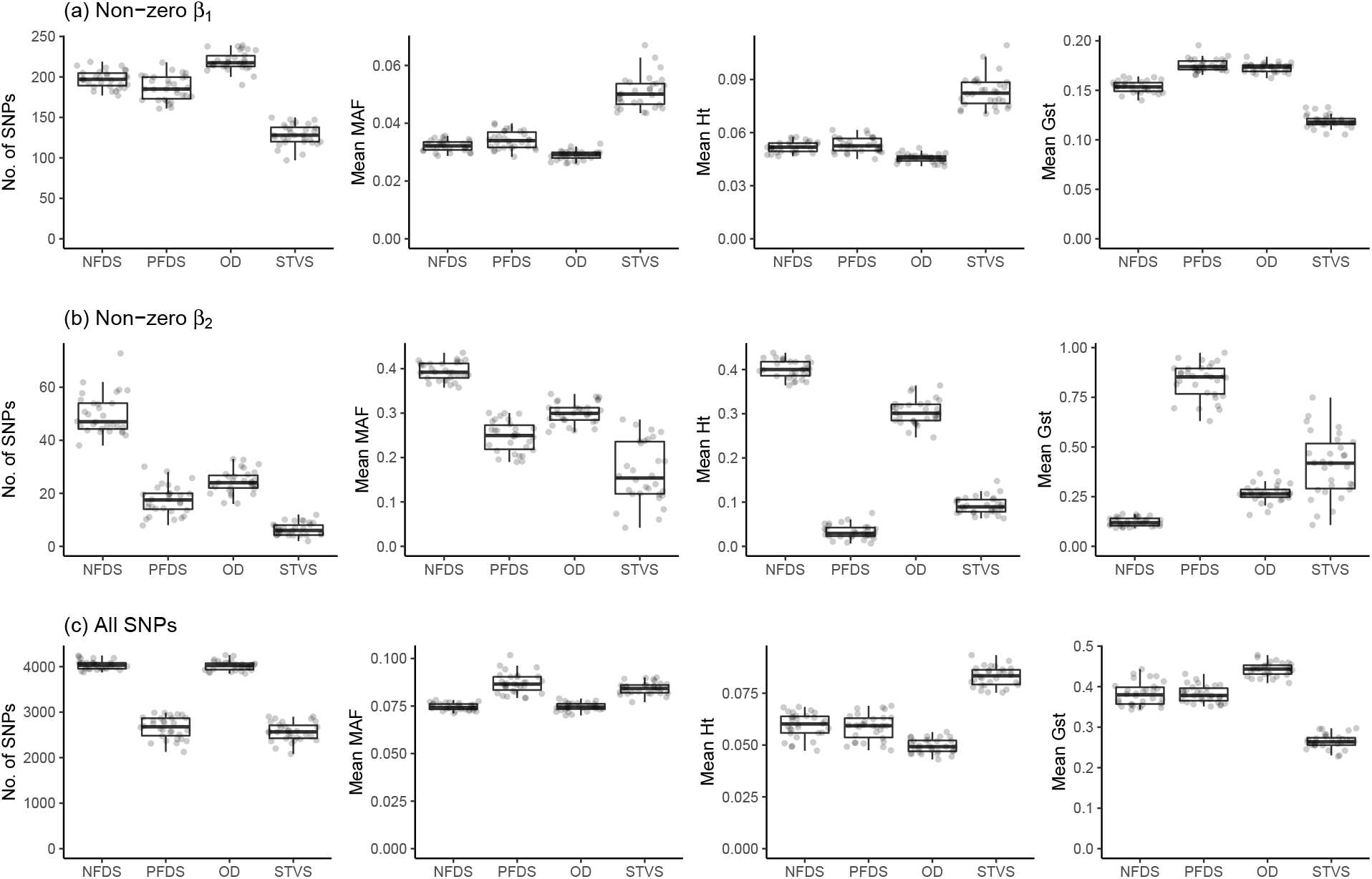
Structure of simulated genomes regarding the loci responsible for stabilizing selection (a), the other forms of selection (b), and genome-wide single nucleotide polymorphisms [SNPs](c). Number of SNPs, mean minor allele frequency (MAF), mean heterozygosity (*H*_t_), and mean fixation indices (*G* _st_) are shown among 30 iterations for four scenarios of selection: NFDS, negative frequency-dependent selection; PFDS, positive frequency-dependent selection; OD, overdominance; STVS, spatiotemporally varying selection.

**Figure S5:**
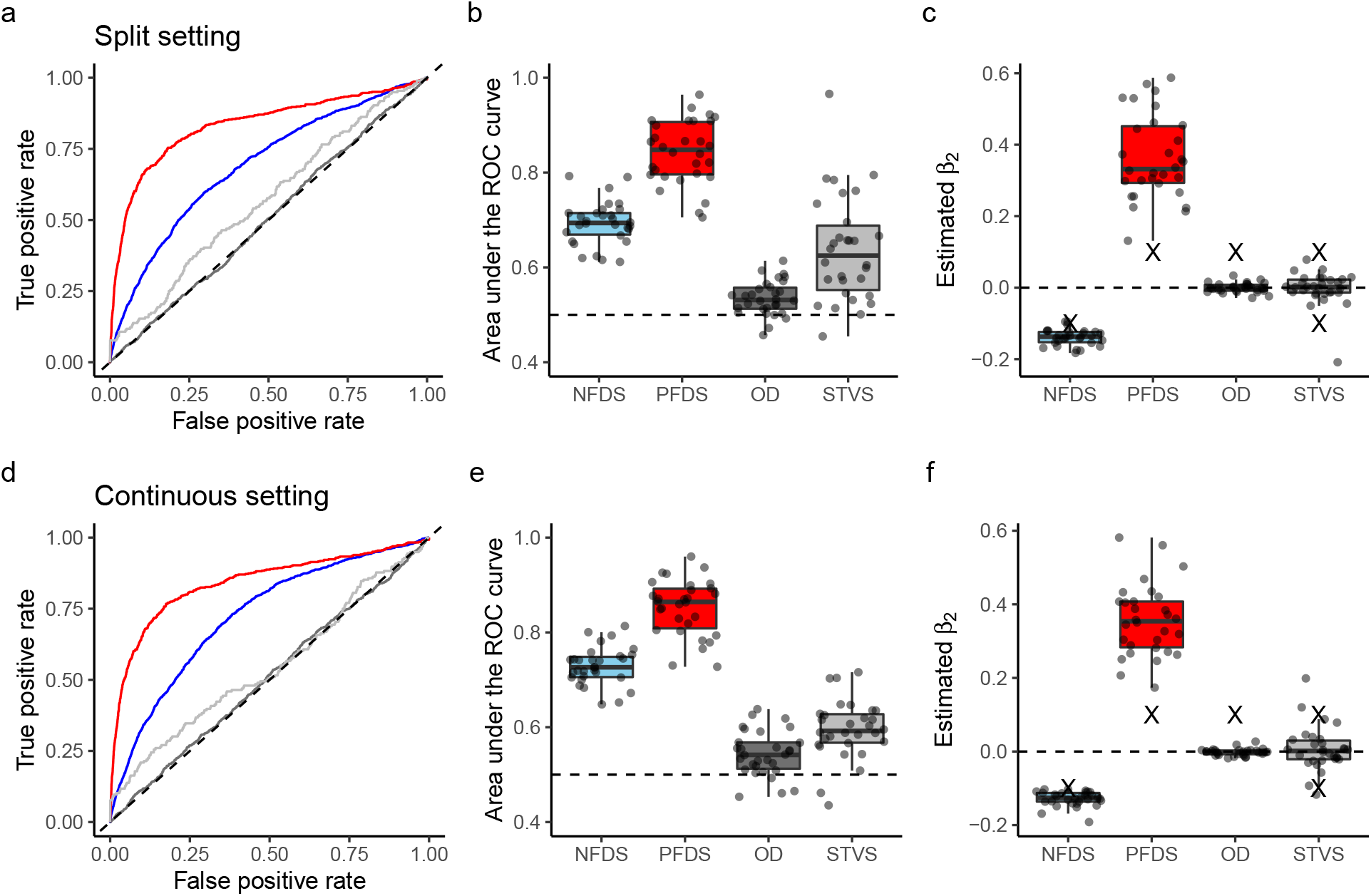
Performance of standard linear models to estimate four types of simulated selection: NFDS, negative frequency-dependent selection; PFDS, positive frequency-dependent selection; OD, overdominance; and STVS, spatiotemporally varying selection. The top and bottom panels show the results of the split and continuous settings, respectively (Fig. 1a). The left panels show the receiver operating characteristic (ROC) curve, which indicates the relationship between the true positive rate and false positive rate. Line colors indicate different simulation scenarios (blue, NFDS; red, PFDS; black, OD; gray, STVS). The middle panel shows the area under the ROC curve (AUC). Dashed lines at AUC = 0.5, indicate no power to detect causal single nucleotide polymorphisms (SNPs). The right panels show the estimated *β*_2_ of causal SNPs, where negative and positive values indicate negative and positive FDS, respectively. Cross marks indicate the true simulated magnitude of *β*_2_.

**Figure S6:**
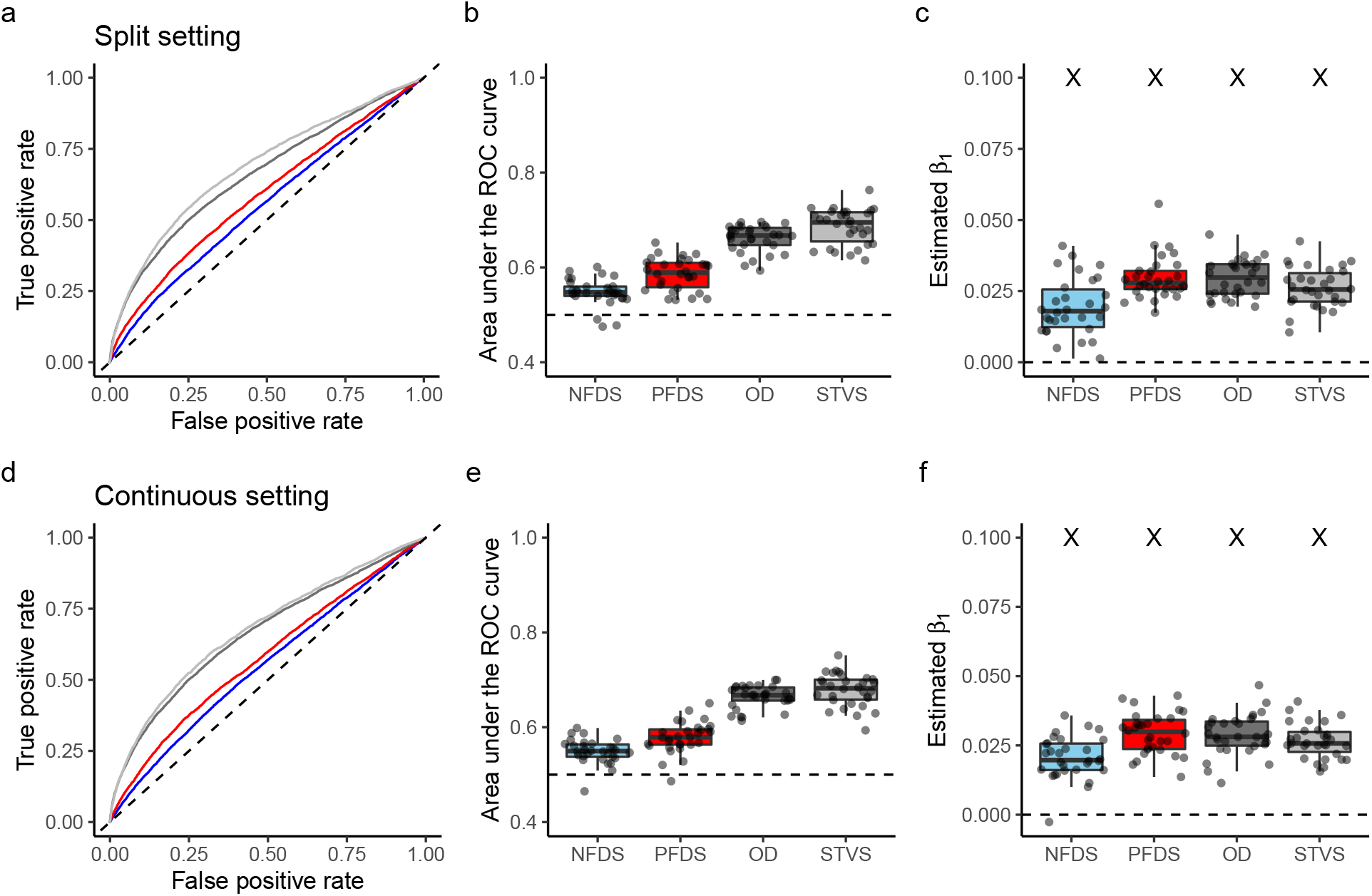
Performance of linear mixed models to detect stabilizing selection under its joint action with four types of selection: NFDS, negative frequency-dependent selection (blue); PFDS, positive frequency-dependent selection (red); OD, overdominance (dark gray); and STVS, spatiotemporally varying selection (light gray). The top and bottom panels show the results of the split and continuous settings, respectively (Fig. 1a). The left panels show the receiver operating characteristic (ROC) curve, which indicates the relationship between the true positive rate and false positive rate. The middle panels show the area under the ROC curve (AUC). Dashed lines at AUC = 0.5, indicate no power to detect causal single nucleotide polymorphisms (SNPs). The right panels show the estimated *β*_1_ of causal SNPs, where negative and positive values indicate negative and positive FDS, respectively. Cross marks indicate the true simulated magnitude of *β*_2_.

**Figure S7:**
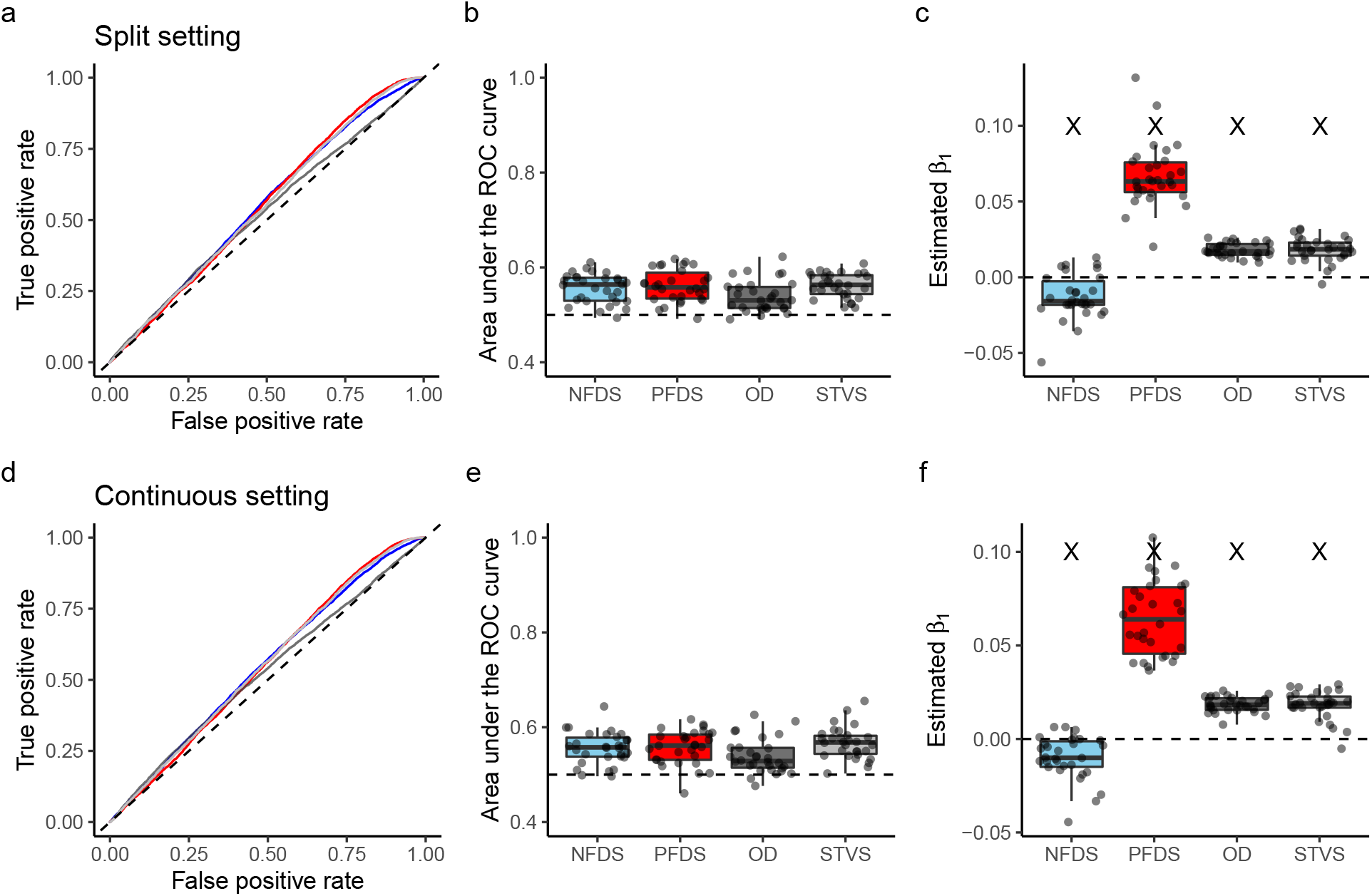
Performance of standard linear models to detect stabilizing selection under its joint action with four types of selection: NFDS, negative frequency-dependent selection (blue); PFDS, positive frequency-dependent selection (red); OD, overdominance (dark gray); and STVS, spatiotemporally varying selection (light gray). The upper and lower panels show the results of the split and continuous settings, respectively (Fig. 1b). The left panels show the receiver operating characteristic (ROC) curve, which indicates the relationship between the true positive rate and false positive rate. The middle panels show the area under the ROC curve (AUC). Dashed lines at AUC = 0.5, indicate no power to detect causal single nucleotide polymorphisms (SNPs). The right panels show the estimated *β*_1_ of causal SNPs, where negative and positive values indicate negative and positive FDS, respectively. Cross marks indicate the true simulated magnitude of *β*_2_.

**Figure S8:**
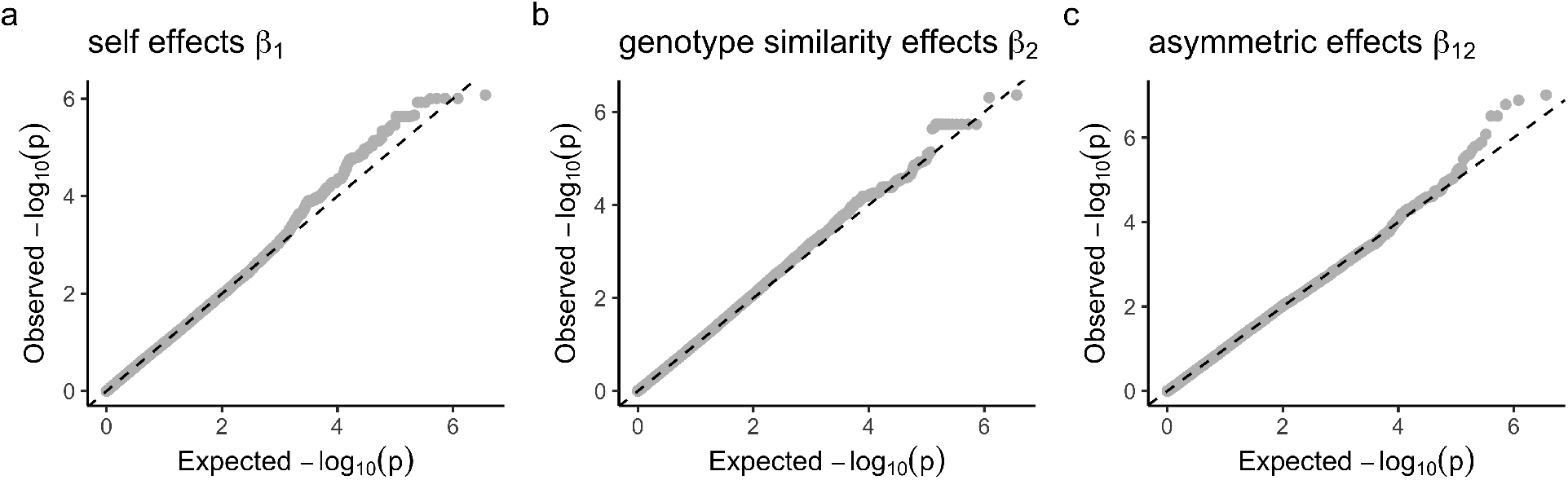
QQ-plots showing observed and expected *p*-values for the genome-wide association studies of branch number in field-grown *A. thaliana*. The results obtained using the linear mixed models are shown. The left (a), middle (b), and right (c) panels display self-genotype, genotype similarity, and asymmetric effects, respectively. Dashed lines indicate the identity between the observed and expected *p*-value scores.

**Figure S9:**
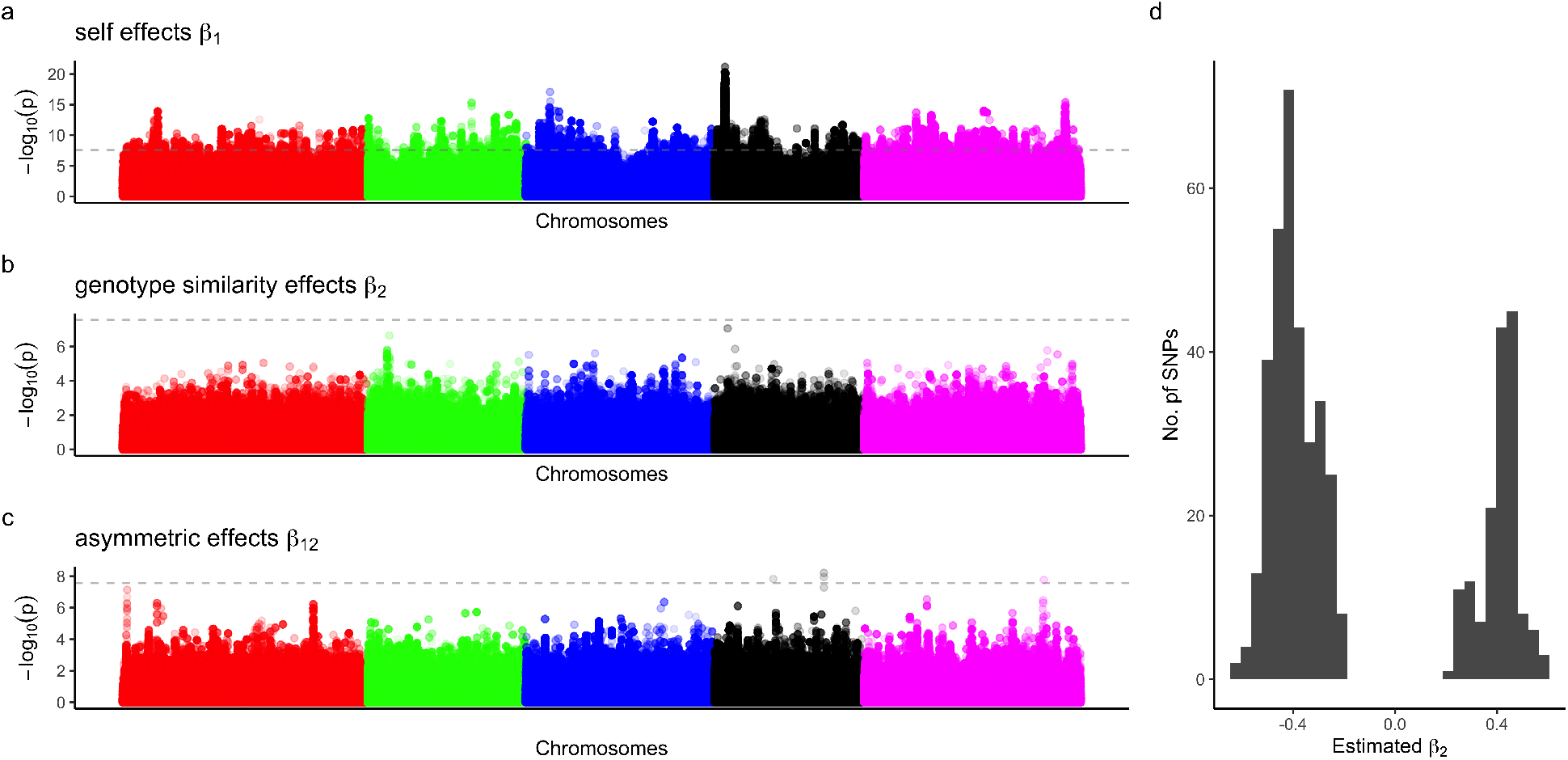
Genome-wide association studies of branch number in field-grown *A. thaliana*. The results of standard linear models are presented. (a, b, and c) Manhattan plots for self-genotype effects, genotype similarity effects, and asymmetric effects, respectively. Horizontal dashed lines indicate *p*-value of *<* 0.05, after Bonferroni correction. (d) Histogram of estimated *β*_2_ among single nucleotide polymorphisms (SNPs) exhibiting *p*-values of *<* 0.0001. Negative and positive *β*_2_ infer loci responsible for negative and positive frequency-dependent selection, respectively.

**Figure S10:**
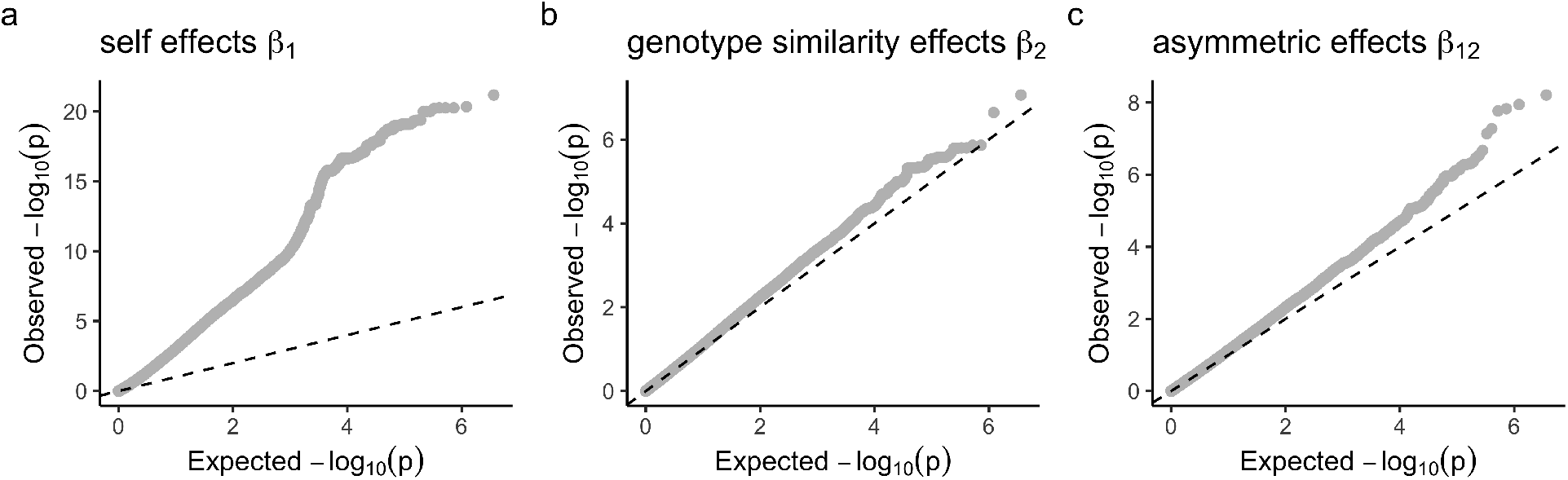
QQ-plots showing observed and expected *p*-values for the genome-wide association studies of branch number in field-grown *A. thaliana*. The results of standard linear models are presented. The left (a), middle (b), and right (c) panels display self-genotype, genotype similarity, and asymmetric effects, respectively. Dashed lines indicate the identity between the observed and expected *p*-value scores.

